# Analysis of Complex DNA Rearrangements During Early Stages of HAC Formation

**DOI:** 10.1101/2020.07.02.184408

**Authors:** Elisa Pesenti, Mikhail Liskovykh, Koei Okazaki, Alessio Mallozzi, Caitlin Reid, Maria Alba Abad, A. Arockia Jeyaprakash, Natalay Kouprina, Vladimir Larionov, Hiroshi Masumoto, William C. Earnshaw

**Affiliations:** Wellcome Trust Centre for Cell Biology, Edinburgh, United Kingdom; National Cancer Institute, National Institutes of Health, Bethesda, United States; Kazusa DNA Research Institute, Kisarazu, Japan

**Keywords:** centromere, Human Artificial Chromosome, Epigenetic Engineering, mitosis, kinetochore, CENP-A

## Abstract

Human Artificial Chromosomes (HACs) are important tools for epigenetic engineering, for measuring chromosome instability (CIN) and possible gene therapy. However, their use in the latter is potentially limited because the input HAC-seeding DNA can undergo an unpredictable series of rearrangements during HAC formation. As a result, after transfection and HAC formation, each cell clone contains a HAC with a unique structure that cannot be precisely predicted from the structure of the HAC-seeding DNA. Although it has been reported that these rearrangements can happen, the timing and mechanism of their formation has yet to be described. Here we synthesized a HAC-seeding DNA with two distinct structural domains and introduced it into HT1080 cells. We characterized a number of HAC-containing clones and subclones to track DNA rearrangements during HAC establishment. We demonstrated that rearrangements can occur early during HAC formation. Subsequently, the established HAC genomic organization is stably maintained across many cell generations. Thus, early stages in HAC formation appear to at least occasionally involve a process of DNA shredding and shuffling that resembles chromothripsis, an important hallmark of many cancer types. Understanding these events during HAC formation has critical implications for future efforts aimed at synthesizing and exploiting synthetic human chromosomes.

## Introduction

Human Artificial Chromosomes (HACs) are non-essential mini-chromosomes that replicate and segregate correctly in human cells^1^. HACs made using synthetic centromeric DNA have been extensively characterized and improved over the last decade^2–5^. They now represent an important tool to study epigenetic regulation of centromere structure and function^2,6–10^, to study full-length gene functions in mutant animal or human cells^5,11–16^, to measure chromosome instability (CIN) and identify new targets for cancer therapy^17–19^. Also, synthetic HACs do not interfere with embryogenesis in mouse, making them a promising tool for future gene therapeutic studies^15^. Synthetic HACs were originally designed using a “bottom up” approach to contain only pre-defined DNA arrays^1,20,21^. Their design allows the HAC centromeres to be easily modified and inactivated/removed by targeting with chimeric proteins specifically directed to the synthetic DNA^2,4,11,14^. However, they also present a number of challenges that must be overcome to enable them to be exploited fully.

HACs form in a complex and as-yet incompletely characterized process after transfection of seeding DNA into human cells. During this relatively lengthy period (typically ∼3 weeks), the HAC-seeding DNA undergoes spontaneous multimerization and in at least some cases may pass through a stage when it is transiently inserted into the arm of an endogenous chromosome^3,22^. The multimerization presumably allows it to attain a threshold size required for stable chromosome segregation^23^. Only the alphoid^tetO^ HAC has previously been characterized in molecular detail. That analysis found complex rearrangements in the organization of its seeding DNA during this multimerization process, including inversions and deletions^3^. These rearrangements are unpredictable and uncontrollable as they occur during the clonal expansion before HAC-bearing cell lines are established and identified. This is the time during which the HAC-seeding DNA is forming a functional centromere, an absolute requirement for the DNA to be stably maintained in cells. As a result, each HAC-bearing cell clone obtained after the selection process contains a HAC that could potentially have a DNA organization different from its sister clones. Although it is known that rearrangements can happen^3^, how and when they occur remains unknown. Importantly, it is not known if these structural rearrangements are an obligate part of the selection process that occurs during centromere formation.

To investigate further how and when rearrangements happen, here we developed a novel alphoid^2domain^ HAC, based on the structure of the alphoid^tetO^ HAC. That first synthetic HAC was constructed from centromeric DNA with a dimeric structure^2^. One monomer of the alphoid^tetO^ HAC was derived from the centromere of chromosome 17 type I *α*-satellite DNA containing a CENP-B box. The other monomer was wholly synthetic alphoid DNA and carried a Tet operator (TetO) in place of the CENP-B box. The CENP-B box is found on all human chromosomes (except the Y) and is a 17 bp sequence recognized by the protein CENP-B^24,25^. This protein’s function is still under investigation, but CENP-B binding is required for stable deposition of the centromeric histone H3-variant CENP-A when HAC-seeding DNA is introduced into cells^26–29^. The presence of the TetO sequence on the synthetic HAC allows the centromeric DNA to be targetable with chimeric Tet repressor (TetR)-fusion proteins that can manipulate the chromatin environment of the centromere and therefore modify the behavior of the HAC centromere.

Here, we have performed the first systematic study of rearrangements that occur during HAC formation and determined how they alter the epigenetic landscape in the HAC centromere and how they impact HAC segregation in mitosis. The seeding DNA of the new alphoid^2domain^ HAC resembles that used to construct the alphoid^hybrid^ HAC described earlier^4^ but was much larger, as we hypothesized that this might minimize the need for rearrangements and amplification during HAC formation. The alphoid^2domain^ HAC contains a CENP-B-containing centrochromatin array and a non-CENP-B-containing domain. The presence of two different domains allows simultaneous targeting of centromeric and flanking regions with different (TetR, LacI and Gal4) fusion proteins and also makes it possible to track rearrangements within and between the arrays. Using this new alphoid^2domain^ HAC, we demonstrate that dramatic DNA rearrangements can occur early during HAC formation and that once formed, they are stably maintained across many cell generations. Thus, a time-limited disruptive event of DNA shredding and shuffling, possibly involving a process resembling chromothripsis^30–32^, can occur early during centromere establishment in human cells.

## Results

### Generation of synthetic *α*21-I^TetO^ and *α*21-II^LacO/Gal4^ arrays using tandem ligation array amplification

The alphoid^2domain^ HAC is formed by two arrays, similarly to a previously constructed alphoid^hybrid^ HAC^4^, but using a ∼2.5x larger (∼120 kb) HAC-seeding construct. The CENP-B-containing centrochromatin array was designed to resemble the previously published alphoid^tetO^ HAC^2^, but in this case, using 11-mer (1886 bp) high order repeats (HORs) of alphoid type I DNA from the centromere in human chromosome 21. Each monomer of this synthetic HOR contains either a 17 bp CENP-B box, essential for CENP-A deposition^29,33^ or a 39 bp tetracycline operator (TetO) targetable sequence, which is the binding site for *E. coli* tetracycline repressor (TetR). This dimer is the basic unit for the so-called *α*21-I^TetO^ (TetO) array, which consists of alternating CENP-B-containing and TetO (non-CENP-B)-containing monomers (Figures 1A, S1A).

**Figure 1:**
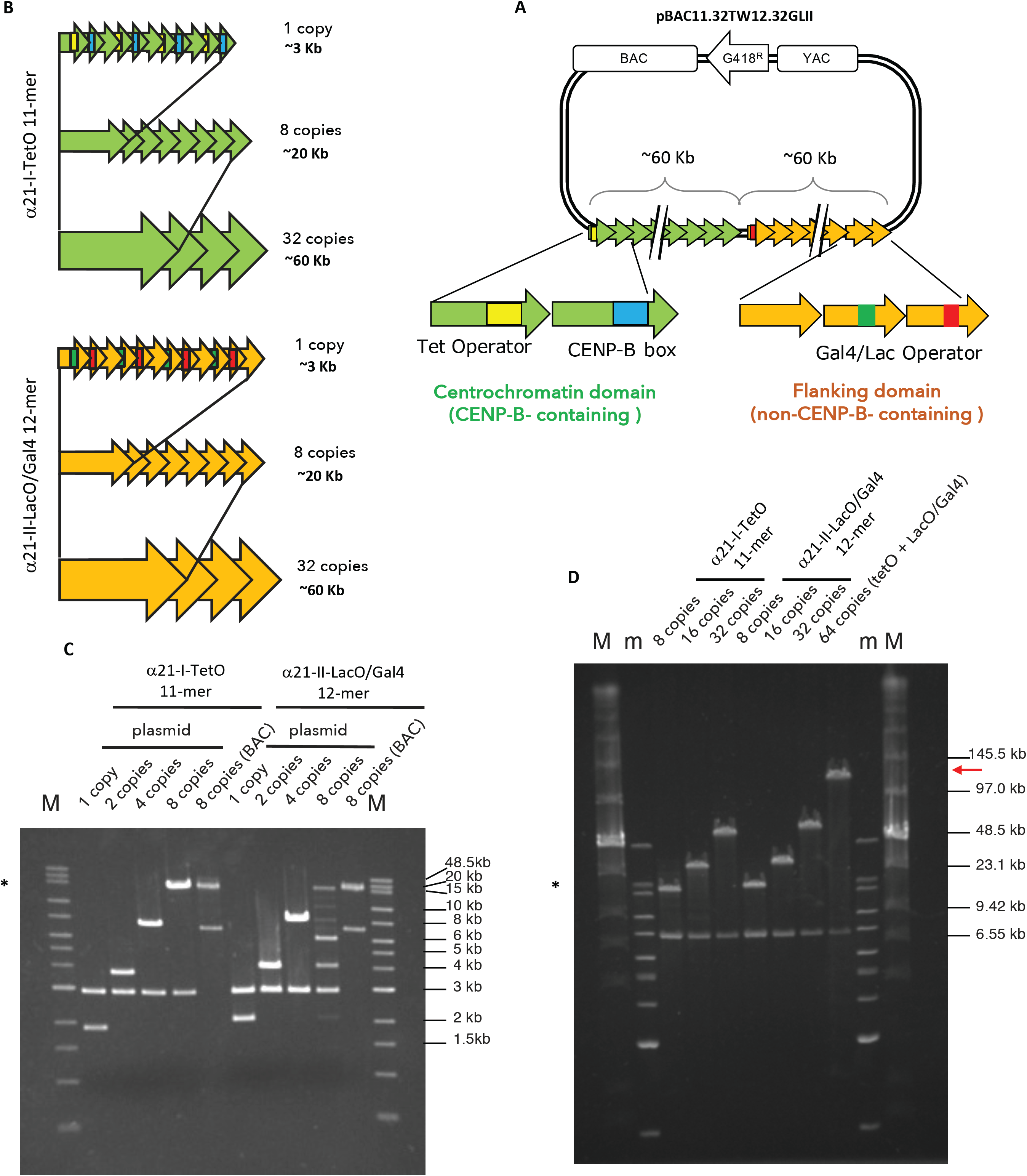
Generation of synthetic *α*21-I^TetO^ and *α*21-II^LacO/Gal4^ arrays. (A) Scheme of the pBAC11.32TW12.32GLII containing BAC and YAC cassettes, G418 resistance cassette and synthetic DNA: *α*21-I^TetO^ formed by high ordered repeats (HOR) monomers (green arrows) containing CENP-B boxes (blue) alternate with monomers containing TetO (yellow); *α*21-II^LacO/Gal4^ formed by high ordered repeats (HOR) monomers (yellow arrows) containing Gal4 binding sequence (green) alternating with LacO (red). (B) Schematic of the assembly of the *α*21-I^TetO^ and *α*21-II^LacO/Gal4^ arrays. (C, D) PFGE analysis of the nascent *α*21-I^TetO^ and *α*21-II^LacO/Gal4^ arrays, cut with BamHI/NotI after each cycles of tandem ligation array amplification as described in Figure S2A (C) and Figure S2B (D). Expected sizes: *α*21-I^TetO^ 11-mer 1 copy (1.9 kb), 8 copies (15.2 kb), 32 copies (60.8 kb); *α*21-II^LacO/Gal4^ 12-mer 1 copy (2 kb), 8 copies (16 kb), 32 copies (64 kb). Plasmid vector is 2.9 kb, BAC vector is 7.1 kb. The asterisk (*) indicates the fragments that have been cloned into BAC vector (8 copies, 16 kb); red arrow in D indicates the size of the final pBAC11.32TW12.32GLII (∼120 kb) (m and M, markers).

The other, non-CENP-B-containing, array is comprised of repeated segments of *α*-satellite type II DNA, lacking CENP-B boxes. In endogenous chromosomes these sequences form the pericentromeric heterochromatin flanking the centromere. To allow targeting of this non CENP-B-containing array with different fusion proteins, LacO and Gal4-targetable sequences were embedded in the array, as previously described^4^. This allows its targeting by chimeric fusions to either *E. coli* lactose repressor (LacI) and/or the yeast Gal4 protein. We refer to this non-CENP-B-containing array as the *α*21-II^LacO/Gal4^ (LacO/Gal4) array (Figure 1A – Figure S1A).

Our initial cloning efforts yielded a *α*21-I^TetO^ 11-mer (1886 bp) and a *α*21-II^LacO/Gal4^ 12-mer (2068 bp) in a plasmid backbone (Figure S1B, S1C). Each basic unit of this 11-mer or 12-mer was then elongated by tandem-ligation-amplification until fragments containing 8 copies were obtained (Figure 1B - Figure S2A). In this tandem-ligation-amplification, cycles of restriction enzyme digestion were performed and followed by ligation as shown in Figure S2A. Upon each cycle of ligation, the restriction site joining the two units was lost, so the next digestion occurred without cutting the nascent elongating array. In this case, cycles of SpeI/ScaI and NheI/ScaI digestions were performed (Figure S2A). After each round of restriction digestion and ligation, the nascent DNA was cut with BamHI and NotI, in order to separate the insert from the 2.9 kb vector, and subsequently analyzed by agarose gel electrophoresis (Figure 1C). Ultimately, the highest molecular weight band (16.6 kb for 8 copies, marked with *) was excised and cloned into a BAC vector capable of more stably maintaining longer repetitive sequences. The structure of the BAC vector is shown in Figure S1D.

Starting from a BAC clone carrying 8 copies of the 11-mer and 12-mer, the tandem-ligation-amplification process was repeated until the inserts reached 32 copies (∼60 kb). To do that, cycles of SpeI/KasI and NheI/KasI digestions were performed (Figure S2B). As before, the nascent array was cut with BamHI and NotI after each reaction of restriction digestion and ligation to separate the insert from the 7.1 kb BAC vector and analyzed by agarose gel electrophoresis (Figure 1D).

Ultimately, the two complete *α*21-I^TetO^ and *α*21-II^LacO/Gal4^ arrays (∼60 kb each) were joined together by tandem ligation into a pBAC vector containing a G148 resistance gene (Figure 1A, D – Figure S2B). As a result, we obtained the HAC-seeding construct pBAC11.32TW12.32GLII, carrying 32 copies of the *α*21-I^TetO^ 11-mer and 32 copies of the *α*21-II^LacO/Gal4^ 12-mer, with a total length of ∼120 kb (Figure 1D, red arrow). This input DNA was then amplified in bacteria prior to transfection into human HT1080 fibrosarcoma cells for HAC formation.

### Isolation of input pBAC11.32TW12.32GLII DNA with equal amounts of *α*21-I^TetO^ and *α*21-II^LacO/Gal4^ repeats

To amplify the HAC-seeding construct, pBAC11.32TW12.32GLII DNA was electroporated into *E. coli* DH10B and the size of the array was determined by CHEF (Contour-clamped Homogeneous Electric Field) gel electrophoresis. In all, 16 BACs isolated from different bacterial clones were obtained from large scale bacterial cultures and digested with NotI and BamHI to release the HAC-seeding array from the BAC backbone (Figure 2A, B). Gel electrophoresis revealed that 8 out of 16 colonies (labeled in red in Figure 2A) maintained the original ∼120 kb length of the synthetic BAC DNA (Figure 2A, red arrow).

**Figure 2:**
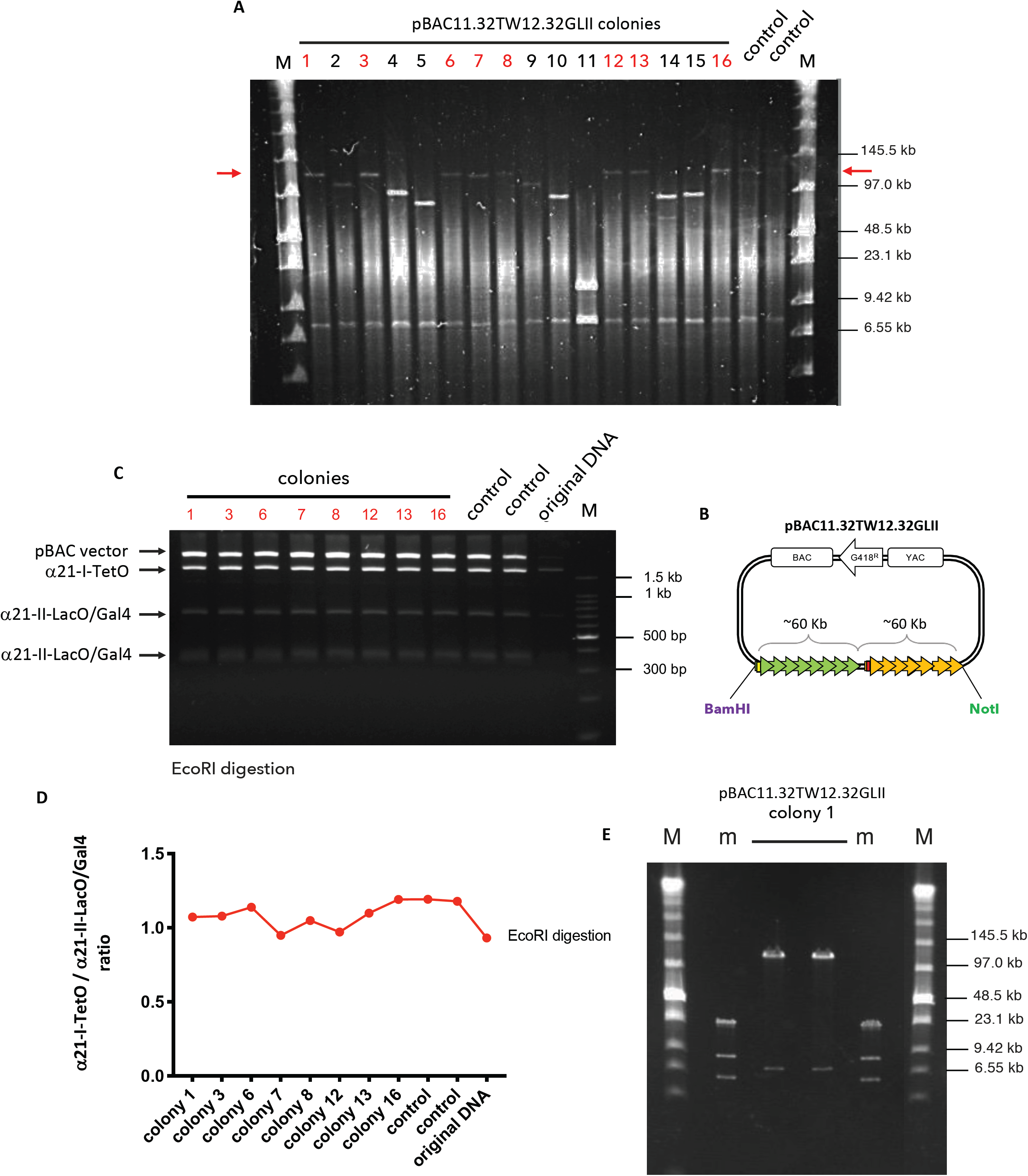
Formation of input pBAC11.32TW12.32GLII DNA. (A) CHEF analysis of 16 bacterial DNA after transformation with pBAC11.32TW12.32GLII and NotI and BamHI digestion: red arrows indicate the size of the final vector (∼120 kb); colonies labelled in red contain the insert of the desired length. DNA used for transfection as a control (in duplicate) (M marker). (B) Scheme of the pBAC11.32TW12.32GLII input DNA showing restriction sites for NotI and BamHI used to release the synthetic DNA. (C) PFGE analysis of selected bacterial colonies (in red) digested with EcoRI: each fragment originates from a different array (label on the left). DNA used for transfection as a control (in duplicate); original DNA as uncut sample (M marker). (D) *α*21-I^TetO^ and *α*21-II^LacO/Gal4^ DNA ratio calculated with ImageJ on the intensity of the bands shown in C for each bacterial colony. Control and original DNA as in C. (E) CHEF analysis of bacterial colony #1 DNA (in duplicate) digested with NotI and BamHI to release the synthetic DNA (m and M, markers).

To establish a HAC with an equal amount of CENP-B-containing and non-CENP-B-containing chromatin, it was important to confirm that the HAC-seeding pBAC11.32TW12.32GLII DNA carries an equal number of *α*21-I^TetO^ and *α*21-II^LacO/Gal4^ arrays. To do so, we digested the input DNA with EcoRI, a restriction enzyme that cuts both the arrays and the BAC vector (Figure S1B, C, D; only single-cut restriction enzymes are shown). We predicted in silico how the EcoRI restriction pattern should look (Figure S1E) and whether each band originates from the *α*21-I^TetO^ or the *α*21-II^LacO/Gal4^ array. The *α*21-I^TetO^ array cut with EcoRI should produce a fragment of 1880 bp, while the *α*21-II^LacO/Gal4^ array should produce fragments of 677, 370, 342, 340 and 339 bp. The vector yields a band of 7499 bp (Figure S1E).

The 8 colonies containing ∼120 kb BAC DNA were digested with EcoRI and the corresponding digests run on an agarose gel (Figure 2C). The results in Figure 2C match the prediction in Figure S1E. Analyzing the intensity of the corresponding bands on the agarose gel in Figure 2C using ImageJ, we scored the ratio between *α*21-I^TetO^ and *α*21-II^LacO/Gal4^ arrays. This confirmed that the BAC DNA contains equal amounts of *α*21-I^TetO^ and *α*21-II^LacO/Gal4^ repeats (Figure 2D). Thus, BAC DNA from clone number 1 (number 1 of Figure 2A and 2C; Figure 2E, sample in duplicate) was chosen as our HAC-seeding input DNA for HAC formation in human cells.

### Screening of HT1080 colonies following transfection with HAC-seeding DNA

HAC formation occurs following transfection of the HAC-seeding DNA into a suitable cell line, and colonies originating from single cells grow under selection. During the process, the input DNA is incorporated into the cell nucleus where it can undergo different fates: it can be integrated into a chromosome arm; it can form an autonomous HAC; the cell population can contain a mixture of both (Figure 3A); or, less frequently, the cells can acquire drug resistance but lose the remainder of the input DNA (not shown)^34^. In order to form a HAC, the input DNA must multimerize to reach a threshold size for a stable chromosome^23^. This step occurs naturally after transfection and it is uncontrollable, leading to different levels of amplification of the input DNA within the cell. As a result, after transfection each single colony contains either a HAC, an integration, or a mixture of the two, with a different degree of amplification of the HAC-seeding DNA^34^.

**Figure 3:**
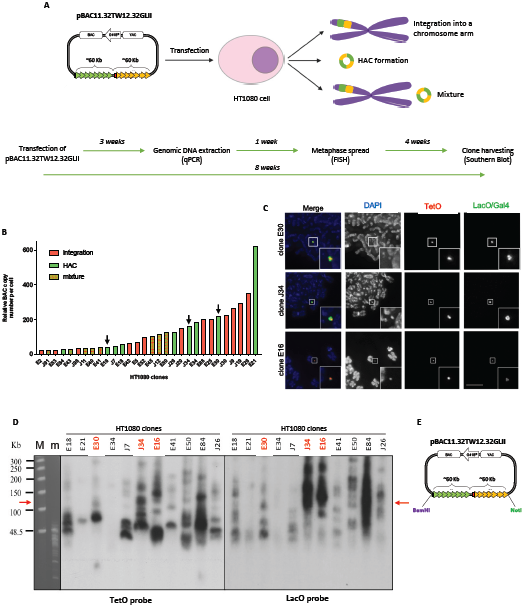
Screening of HT1080 colonies after transfection with pBAC11.32TW12.32GLII. (A) Scheme showing the possible fates of the pBAC11.32TW12.32GLII HAC seeding DNA after transfection in HT1080: in yellow and green (as integration or HAC) is represented the synthetic DNA. Timeline of the experiments performed from transfection into HT1080 cells. (B) BAC copy number (y axis) analyzed by qPCR in each HT1080 clone (x axis): only HT1080 clones containing >20 BAC copies are represented in the graph. HT1080 clones are represented in green (HAC), red (integration) or mixture (both) according to the results of the FISH screening, as shown in C. Black arrows indicate the clones shown in C and analyzed further. (C) Representative pictures of oligo-FISH staining of HT1080 clones: slides have been hybridized with DNA probes (TetO-dig/rhodamine -dig antibody, Gal4-biotin and LacO-biotin/Fitc-streptavidin). DAPI stains DNA. Scalebar = 10µm. (D) Southern blot of selected HT1080 clonal DNA (as labelled on top of the panel) digested with BamHI and separated by CHEF; the transferred membrane was hybridized with radioactively labelled TetO (left) or LacO (right) specific probes. Red arrows indicate the expected size of the band without rearrangements. Clones labelled in red have been screened further (M and m, markers). (E) Cartoon of the pBAC11.32TW12.32GLII input DNA showing restriction sites for NotI and BamHI.

As previously published^2,4^, we chose to transfect pBAC11.32TW12.32GLII into HT1080 cells. This fibrosarcoma cell line has a chromatin state permissive for HAC formation due to having a relatively low level of H3K9me3 as a result of decreased expression of Suv39h1 methyltransferase^10^. In cells with higher Suv39h1 expression, CENP-A assembles on HAC-seeding DNA, but is subsequently displaced by invading H3K9me3-containing heterochromatin^10^.

pBAC11.32TW12.32GLII from clone 1 (Figure 2E) was transfected into HT1080 cells and single cell clones were grown for 3 weeks in media containing geneticin. We collected genomic DNA from 124 resistant colonies and measured the BAC copy number by qPCR to obtain an approximate measurement of the degree of amplification of HAC-seeding DNA. Primers specific for the alphoid^2domain^ HAC were designed and a different HAC with a known BAC copy number was used as standard.

30 HT1080 colonies containing detectable amounts of HAC-seeding BAC sequences (copy numbers >20), were then screened by fluorescent in situ hybridization (FISH) for the presence of HACs, integrations, or mixtures of both (Figure 3B). FISH to detect pBAC11.32TW12.32GLII was performed 4 weeks after transfection (timeline in Figure 3) using TetO and LacO-specific oligos labelled with fluorochromes (see Methods for details). Figure 3B presents data of the qPCR analysis combined with the results of the FISH screening. Representative images from the FISH screening of selected HT1080 clones are shown in Figure 3C. HACs can be visualized as discrete spots by DAPI staining (Figure 3C). Interestingly, the size of each HAC estimated by FISH correlated the results of qPCR, with the larger HACs corresponding to higher BAC copy numbers and vice versa (Figure 3B black arrows, C). For simplicity, we discarded HAC-containing clones with more than 1 HAC.

In all HACs, the *α*21-I^TetO^ array seems to localize to the center of the HAC, where it is surrounded by the *α*21-II^LacO/Gal4^ arrays (Figure 3C). This organization corresponds to that seen for previously published HACs^4^, and presumably reflects the HAC structure, with a CENP-B-containing centromere surrounded by pericentromeric heterochromatin.

### pBAC11.32TW12.32GLII forms HACs more efficiently than previous HAC-seeding constructs

In order to determine the fate of the HAC-seeding DNA, a minimum of 25 metaphases were screened by FISH for each clone. In the screening shown in Figure 3B, 30% (9/30) of HT1080 colonies contained only HACs, 43% (13/30) contained integrations, and 26.6% (8/30) contained a mixture of both. This frequency of HAC-containing colonies for the alphoid^2domain^ HAC is ∼3 times higher than in previous studies^2,4,33^. The efficiency of HAC formation by pBAC11.32TW12.32GLII was comparable to the frequency of HAC formation reported in our previous study when cells were stably co-transfected with the HAC seeding DNA plus CENP-A directed to the synthetic centromere^4^. It is possible that using a larger HAC-seeding DNA may spontaneously increase the efficiency of CENP-A deposition on the centromeric DNA.

### Analysis of rearrangements of the pBAC11.32TW12.32GLII arrays in HAC-containing HT1080 colonies

We wished to determine whether the size amplification that occurred during early stages of alphoid^2domain^ HAC formation was also accompanied by rearrangements of the HAC-seeding DNA arrays. To perform a structural analysis of the *α*21-I^TetO^ and *α*21-II^LacO/Gal4^ arrays, we performed Southern blot analysis using TetO and LacO specific probes (Figure 3D). Cell clones were grown for 8 weeks, or approximately 50 doublings of the HT1080 cells, prior to Southern blot analysis (timeline in Figure 3). Genomic DNA from 9 HAC-containing cell lines (and 2 two HT1080 clones containing integrations as control) was digested with BamHI, which has a unique site only on the vector backbone (Figure 3E). The DNA fragments were separated by CHEF gel electrophoresis and the membranes hybridized with TetO and LacO-specific probes.

If only simple multimerization of pBAC11.32TW12.32GLII occurred during HAC formation, the Southern blots should display a single ∼120 kb band, corresponding to the size of digested input DNA. Importantly, since the restriction enzyme cuts only at the edge of the *α*-satellite arrays, this should be the case regardless of whether the Southern blot analysis uses the TetO or LacO probe. Surprisingly, none of the analyzed clones showed this single ∼120 kb band (Figure 3D, red arrow). Instead, each clone has a different number of DNA fragments of different sizes, and these also vary for each clone for the two probes. Many bands are smaller than the 120 kb input band, but some are considerably larger. Thus, the arrays of the HAC-seeding pBAC11.32TW12.32GLII DNA underwent a complex series of rearrangements during HAC formation, as described for the alphoid^tetO^ HAC^3^.

Based on the results displayed in Figure 3, we decided to further characterize three clones (E30, J34 and E16) that showed different levels of amplification by qPCR (black arrows in Figure 3B) and different numbers and sizes of rearrangements by Southern blotting (labelled in red in Figure 3D).

### pBAC11.32TW12.32GLII undergoes multiple rearrangements during early stages of HAC formation

The rearrangement of HAC-seeding DNA was previously described for the single-domain alphoid^tetO^ HAC^3^. It was proposed that the HAC-seeding DNA structure may continue to change and evolve for weeks or possibly months after HAC transfection. This raises the possibility that the populations analyzed in Figure 3D might consist of mixtures of alphoid^2domain^ HACs with different structures. To test this hypothesis, the three clones E30, J34 and E16 were further subcloned to obtain homogeneous cell populations (timeline in Figure 4; 9 weeks or approximately 55 population doublings after transfection). Initially clone E21 was also subcloned but, unfortunately, we could not grow any subclone with stable HAC segregation, so E21 was excluded from the subsequent analysis.

**Figure 4:**
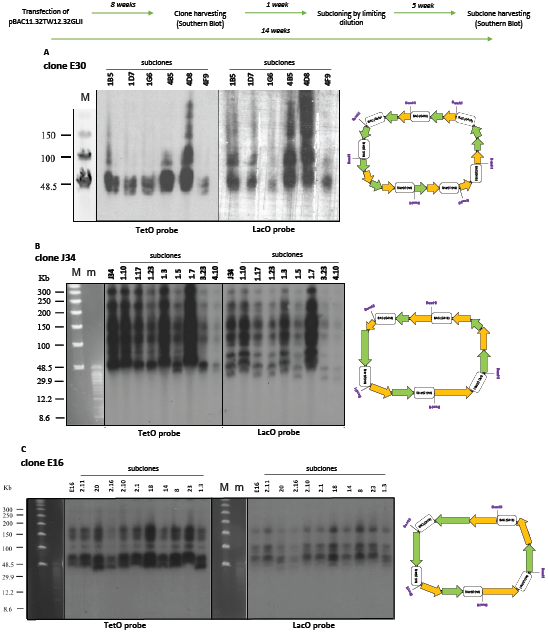
Analysis of rearrangements of the pBAC11.32TW12.32GLII arrays. Timeline of the experiments performed from transfection into HT1080 clones and subclones. (A,B,C) Southern blot of subclones from clone E30 (A), J34 (B) and E16 (C): DNA was digested with BamHI and separated by CHEF. The transferred membrane was hybridized with radioactively labelled TetO (left) or LacO (right) specific probes. Cartoons on the right represent the outcome of rearrangements in the corresponding clone: green arrows represent *α*21-I^TetO^ array, yellow arrows represent *α*21-II^LacO/Gal4^ array, boxes represent the BAC vector (M and m, markers).

Alphoid^2domain^ HAC subclones were isolated by limiting dilution and screened by FISH for the presence and the number of the HACs in each clonal cell line (12 subclones for E30, 35 subclones for J34 and 25 subclones for E16; detailed timeline in Figure S3). Subclones with highly mis-segregating HACs were discarded and subclones with a higher percentage of single HACs per cell were studied further (6 subclones for E30, 8 subclones for J34 and 10 subclones for E16; Figure 4). As an example of the screening, Figure S3A shows representative images from FISH screening of 6 subclones from clone E30. Figure S3B shows the number of metaphases containing 0, 1 or 2 HACs in subclones from clone E30, with percentages indicating the cells with 1 HAC.

To study rearrangements in subsequent cell generations, genomic DNA from the selected subclones originating from E30, J34 and E16 was digested with BamHI and separated by CHEF gel electrophoresis (Figure 4). Surprisingly, Southern blot analysis revealed that all subclones were almost identical in the number and sizes of rearrangements. Furthermore, they all recapitulated the pattern of rearrangements seen in the original clone (Figure 4A-C). As for the original clones, hybridization with the TetO-specific probe consistently yielded a different hybridization pattern from that seen with the LacO-specific probe on the same sample.

Taken together, these data show that the *α*21-I^TetO^ and *α*21-II^LacO/Gal4^ arrays of HAC-seeding DNA pBAC11.32TW12.32GLII independently undergo unique fragmentation, recombination and amplification events during the first 8 weeks of alphoid^2domain^ HAC formation. Subsequently, these rearrangements appear to be maintained stably through cell generations, up to 14 weeks. This agrees with the observation that the HAC structure seems to be stable through multiple cycles of MMCT (Microcell Mediated Chromosome Transfer) in different cell lines^3,35^.

### Different HAC-containing clones show different degrees of rearrangements

Not all alphoid^2domain^ HAC clones analyzed underwent the same degrees of rearrangements. For example, subclones from clone E30 (Figure 4A) display a predominant band around 50 kb in all subclones, and only 2 of the 6 subclones clearly show fragments around 80-100 kb with both probes (subclones 4B5 and 4D8). In contrast, the LacO probe shows 3-4 bands with various signal strengths. This could reflect dimerization/multimerization of the 50 kb fragment observed in the blot. Thus, compared with the other clones analyzed, E30 seems to be less scrambled, with all fragments showing the same size (cartoon in Figure 4A). It therefore appears that during E30 HAC formation pBAC11.32TW12.32GLII underwent an early event in which both the *α*21-I^TetO^ and *α*21-II^LacO/Gal4^ arrays were shortened to roughly 40% of their initial lengths (easiest to imagine if a single deletion of the 120 kb construct occurred spanning the junction between the two arrays), but then were amplified while avoiding further rearrangements.

In marked contrast, clones J34 and E16 displayed a much larger number and variety of rearrangements, with fragments ranging from ∼50 kb up to ∼300 kb (Figure 4B, C and relative diagrams on the right). One possible explanation for this structure is that early during formation of those two HACs and following some initial amplification of the arrays, the nascent HAC-seeding DNA experienced multiple chromosome breaks and shuffling followed by re-ligation, leading to fragments of different sizes.

Interestingly, the smaller array size in clone E30 (Figure 4A) correlates with its larger number of BAC copies (presumably resulting from a larger number of amplification cycles) quantitated by qPCR (Figure 3B). In contrast, J34 and E16 with more complex rearrangements, including those producing much larger fragments (Figure 4B, C), have lower BAC copy numbers (Figure 3B). These observations suggest that the rearrangements may have occurred very early, prior to completion of the multimerization that allowed the HAC to pass the minimum size threshold required to form a stable centromere/kinetochore^23^.

These data show that, as previously suggested^3^, during alphoid^2domain^ HAC formation, the predicted regular structure of the HAC-seeding DNA is disrupted by complex rearrangements whose mechanism remains unknown.

### Visualization of *α*21-I^TetO^ and *α*21-II^LacO/Gal4^ arrays on chromatin fibers

We performed immuno-fiber-FISH analysis to confirm the presence of *α*21-I^TetO^ and *α*21-II^LacO/Gal4^ arrays on stretched DNA fibers, and to also determine the distribution of CENP-A and H3K9me3 on those fibers.

For each set of subclones, one was selected for further experiments, based on the percentage of cells bearing a single HAC (E30 subclone 1B5, J34 subclone 1.10 and E16 subclone 23; as example for clone E30, see Figure S3B). DNA fibers were prepared from the selected subclones and incubated with purified TetR-eYFP or LacI-eYFP fusion proteins expressed in *E. coli* to visualize the corresponding array (TetR-eYFP and LacI-eYFP expressed *in vivo*, both dissociate from the chromatin during fiber preparation) (Figure 5A – Figure S5). Staining of both arrays simultaneously was not possible, since the purified proteins were both tagged with GFP. Attempts to specifically stain fibers with mCherry-TetR isolated from *E. coli* were not successful. Fibers were also stained using CENP-A or H3K9me3-specific antibodies (Figure 5A).

**Figure 5:**
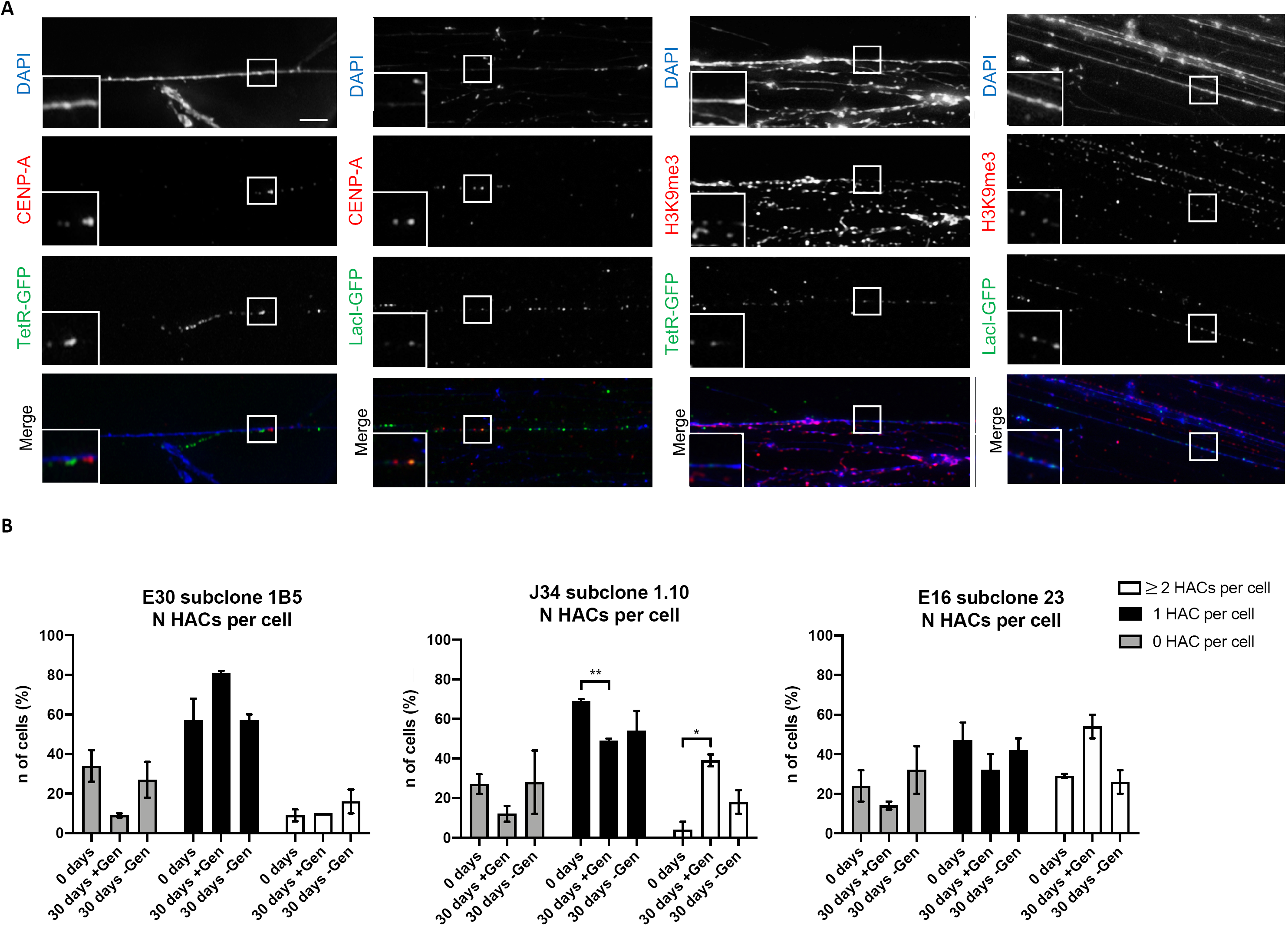
Visualization of *α*21-I^TetO^ and *α*21-II^LacO/Gal4^ arrays on chromatin fibers and mitotic stability of the HAC. (A) Representative pictures of immune-fiber-FISH staining of subclone J34 1.10: slides have been incubated with TetR-eYFP and LacI-eYFP expressed in *E*.*coli* and stained with *α*-CENP-A mouse or *α*-H3K9me3 mouse/TRITC *α*-mouse antibody. DAPI stains the DNA. (B) Number of metaphases (%) containing 0, 1 or *≥*2 HACs for subclones E30 1B5, J34 1.10 and E16 23, after spreading metaphases and hybridizing with TetO and LacO/Gal4 specific probes; total of 2 biological repeats, 50 metaphases for each condition were analyzed. +/-Gen indicates treatment for 30 days from day 0 with (+) or without (-) geneticin. Error bars denote SEM. Statistical test: unpaired t-test (*P < 0.05, **P < 0.01).

Immuno-fiber-FISH experiments revealed that the *α*21-I^TetO^ and *α*21-II^LacO/Gal4^ arrays are both present along these stretched DNA fibers. CENP-A and H3K9me3 are adjacent to both arrays, with no apparent preference for one or the other array (representative images for J34 subclone 1.10 are shown in Figure 5A). The presence of CENP-A and H3K9me3 in close proximity to both arrays can be explained if the rearrangements during alphoid^2domain^ HAC formation lead to a “scrambled” structure of the *α*21-I^TetO^ and *α*21-II^LacO/Gal4^ arrays.

### Geneticin selection enriches the number of HACs in the cell population

To characterize how the mitotic stability of the alphoid^2domain^ HAC is affected by its structure, we performed a stability assay, counting the number of cells with different number of HACs over a period of 30 days with (+) and without (-) geneticin (∼ 25 cell divisions). Metaphase chromosome spreads from E30 subclone 1B5, J34 subclone 1.10 and E16 subclone 23 were analyzed by FISH and imaged at each timepoint using labelled oligos specific for *α*21-I^TetO^ and *α*21-II^LacO/Gal4^, to count cells containing 0, 1 or *≥*2 HACs.

At timepoint zero, each subclone had a characteristic number of cells with 0, 1 or *≥*2 HACs (Figure 5B). Interestingly, 7 days after thawing and while growing in presence of geneticin (corresponding to timepoint zero), the number of cells containing no HAC was higher than expected (20-40% of the cell population, grey bars) for all three subclones. It is possible that after the stress of freezing/thawing, cells were not yet fully responding to the selection.

Importantly, all of the alphoid^2domain^ HACs are extremely stable after 30 days growth in the absence of geneticin, as shown by the number of cells bearing 1 HAC (black bars; daily loss rate of the HAC (R) = 4.76 (± 0.6) x 10^−3^ for E30 subclone 1B5; 8.69 (± 6.67) x 10^−3^ for J34 subclone 1.10; and 3.47 (± 1.66) x 10^−3^ for E16 subclone 23 – see Methods). These measured loss rates were comparable to those measured for previous HACs^2,4^. Thus, alphoid^2domain^ HACs with dramatically different rearrangements of their seeding DNA are very similar in their ability to replicate and segregate correctly during the great majority of cell divisions.

Surprisingly, when cells grow for 30 days (+) geneticin, they seem to acquire a selective advantage for increasing the HAC copy number, as shown by the number of cells bearing *≥*2 HACs (white bars). The accumulation of HAC was particularly evident in J34 subclone 1.10 and E16 subclone 23, while E30 subclone 1B5 did not exhibit this increase (Figure 5B). Notably, the enrichment of cells with 1 or *≥*2 HACs after 30 days (+) geneticin was coupled for all subclones with a reduction in the number of cells with 0 HACs.

The enrichment in cells with *≥*2 HACs could be explained if heterochromatin spreading silences the geneticin resistance gene. In this case, geneticin would select for cells in the population with an increased HAC copy number. Despite the small sample size, it is interesting to note that the alphoid^2domain^ HACs with the more rearranged arrays (J34 1.10 and E16 23) were those where the copy number increased under selection, possibly indicating that the chromatin state is less stable. In contrast, the clone with the least rearranged structure (E30) showed the highest chromatin stability. Taken together these data suggest that the HAC DNA structure may have an impact on HAC chromatin stability over time.

### CENP-A accumulates preferentially on the *α*21-I^TetO^ array with CENP-B boxes

The immuno-fiber-FISH in Figure 5 shows CENP-A and H3K9me3 apparently localized on both the *α*21-I^TetO^ and *α*21-II^LacO/Gal4^ arrays. To better characterize the chromatin state of the two arrays on the HAC-seeding DNA, we performed chromatin immunoprecipitation (ChIP) for CENP-A and several indicative histone modifications using a set of well-characterized monoclonal antibodies^36^, followed by quantitative PCR (ChIP-qPCR) on genomic DNA from E30 subclone 1B5, J34 subclone 1.10 and E16 subclone 23 (scheme of the primers used for qPCR is presented in Figure 6D).

**Figure 6:**
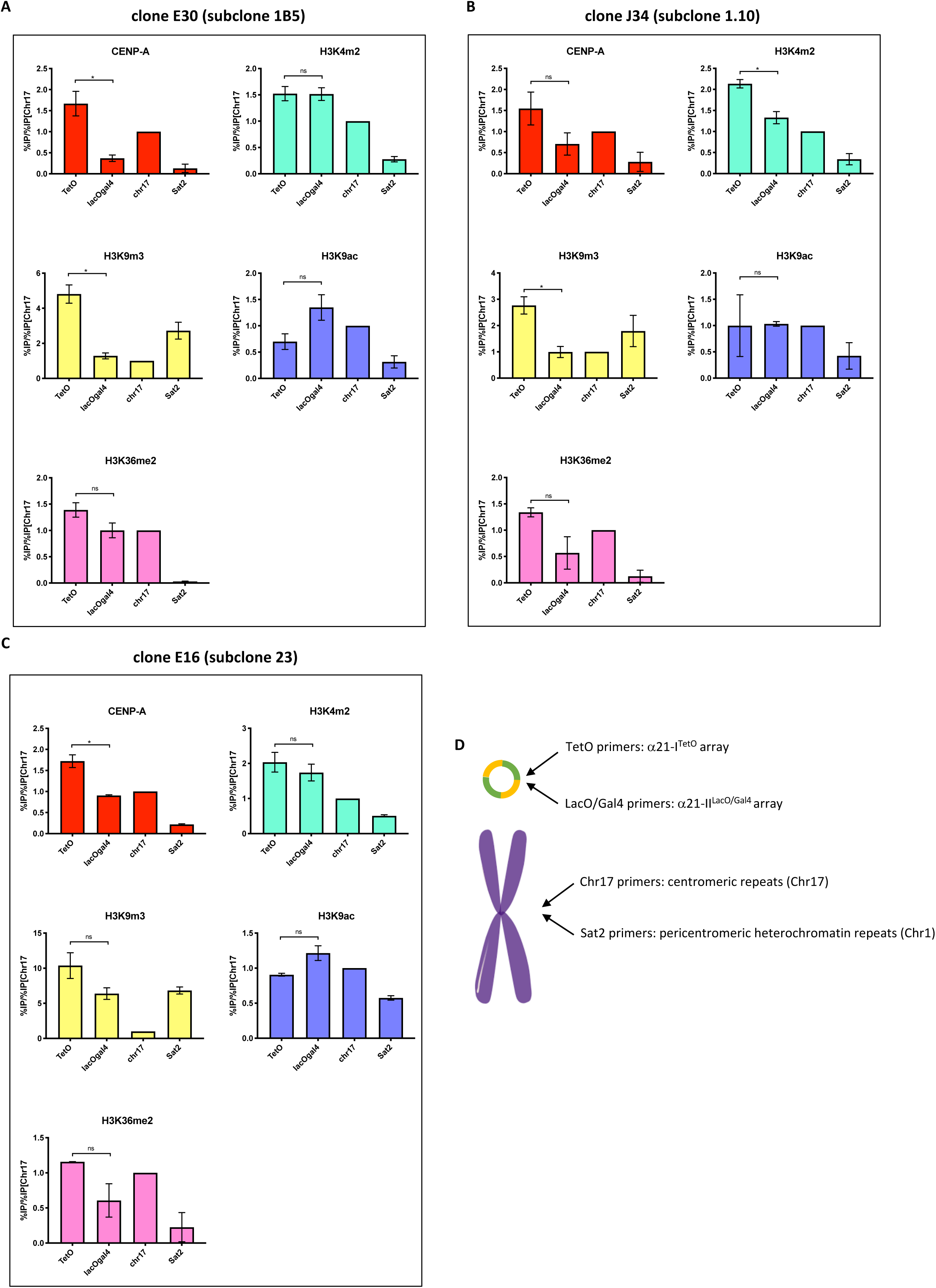
CENP-A accumulates preferentially on the *α*21-I^TetO^ array. (A, B, C) ChIP-qPCR analysis of CENP-A and indicated histone marks modifications in HT1080 subclones E30 1B5 (A), J34 1.10 (B) and E16 23 (C). The *α*21-I^TetO^ array (TetO), *α*21-II^LacO/Gal4^ array (LacOGal4), satellite D17Z1 (chr17) and degenerate satellite II (sat2) repeats were assessed. Values have been normalized against satellite D17Z1 (chr17). Total of 2 biological repeats, n≈5×10^6^ cells each. Error bars denote SEM. Statistical test: Mann-Whitney test (*P < 0.05). (D) Cartoon showing the localization of primers used in qPCR on the HAC and on endogenous corresponding chromosomes.

ChIP data were highly reproducible for the three subclones of the alphoid^2domain^ HAC (Figure 6A, B, C). Thus, the over-all chromatin organization was maintained despite differences in the level of rearrangements. CENP-A accumulated on the *α*21-I^TetO^ array ∼2-3 times more than on the *α*21-II^LacO/Gal4^ array in all the three subclones. This is an average of ∼1.5 times more than on the endogenous centromere of chromosome 17, used as a control (Figure 6A, B, C). This contrasts with a previous study in which the alphoid^hybrid^ HAC was apparently unable to maintain CENP-A only on the centromeric array^4^. It is possible that CENP-A deposition on the *α*21-I^TetO^ array may be favored by the larger size of the input pBAC11.32TW12.32GLII HAC-seeding DNA. CENP-A deposition also correlated with higher levels of H3K4me2 and H3K36me2, as expected for centrochromatin^7,9^. The *α*21-I^TetO^ array also contained a relatively high level of H3K9me3 and a low level of H3K9ac (Figure 6A, B, C), revealing differences from the alphoid^tetO^ HAC, which contained a single HAC-seeding array^2,7^.

Unexpectedly, the *α*21-II^LacO/Gal4^ array had levels of H3K9ac and H3K4me2 (markers for actively transcribed chromatin) ∼ twice those of the satellite II DNA used as a control. Consistent with this observation, levels of H3K9me3 on the *α*21-II^LacO/Gal4^ array were ∼2-4 times lower than on the *α*21-I^TetO^ array (Figure 6A, B, C), revealing a generally open conformation of the chromatin. This was surprising, as we had initially expected this array, which lacks CENP-B boxes, to form heterochromatin. However, given the intermixing of sequences on the HAC-seeding DNA, the presence of strong heterochromatin might have been counter-selected due to its potentially harmful effects on expression of the geneticin-resistance gene.

Taken together these data suggest that *α*21-I^TetO^ array recruits CENP-A and establishes a functional centromere in the alphoid^2domain^ HAC, despite sustaining high levels of H3K9me3.

### *α*21-I^TetO^ and *α*21-II^LacO/Gal4^ arrays do not form functionally independent chromatin domains

To determine whether the molecular structure of the HAC impacts the function of the *α*21-I^TetO^ and *α*21-II^LacO/Gal4^ arrays, we asked whether the two arrays are functionally distinct. To do this, we transiently expressed KAP1 as a chimeric fusion to either TetR-eYFP or LacI-GFP. KAP1 is a scaffolding protein that recruits the CoREST complex, promoting a silent chromatin state and increasing the level of H3K9me3^37^. Previous studies revealed that KAP1 recruitment into the centromere causes a loss of CENP-A and inactivates the kinetochore^38^. Thus, if the two arrays on the alphoid^2domain^ HAC are functionally independent, KAP1 recruitment should have an effect of the HAC centromere only when targeted to the *α*21-I^TetO^ array.

We performed quantitative fluorescent analysis to measure the level of CENP-A and H3K9me3 on the alphoid^2domain^ HAC after targeting KAP1-eYFP fusions to the two arrays both separately and simultaneously for 48h, using the eYFP to localize the HAC arrays in interphase cells (Figure S4A, B). Targeting KAP1 to the *α*21-I^TetO^ array led to a significant (∼2 fold) decrease in CENP-A levels on the HAC for all three subclones analyzed (Figure 7A). This is similar to what was reported for the alphoid^tetO^ HAC^6^. The decrease in CENP-A was accompanied by an increase in H3K9me3 levels when targeting KAP1 to the *α*21-I^TetO^ array (Figure 7B). Different subclones showed different levels of H3K9me3 enrichment, possibly due to intrinsic variation in the H3K9me3 basal levels in each subclone (Figure 6A, B, C).

**Figure 7:**
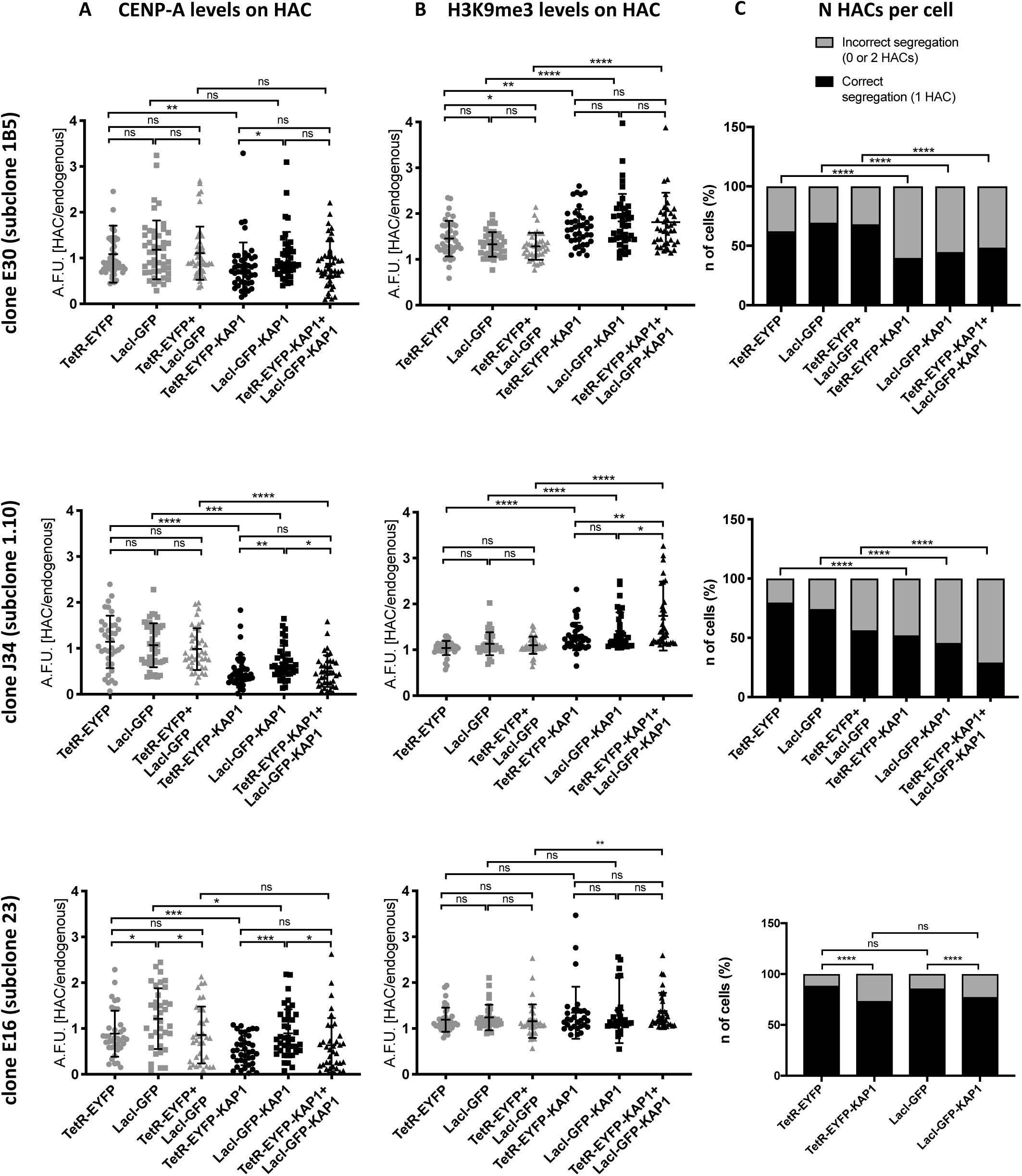
*α*21-I^TetO^ and *α*21-II^LacO/Gal4^ arrays are not functionally independent. (A, B) Quantification of HAC-associated CENP-A staining (A) and H3K9me3 staining (B) in individual cells of each indicated HT1080 subclone 48h after transfection with the indicated fusion proteins; values plotted as A.F.U. (arbitrary forming units). Solid bars indicate the medians, and error bars represent the SD. n = two independent experiments for each staining; ∼30 cells analyzed in each experiment. Asterisks indicate a significant difference (*P < 0.05, **P < 0.01, ***P < 0.001, ****P < 0.0001; Mann–Whitney test). (C) Number of interphase nuclei (%) showing correct segregation (1 HAC) or missegregation (0 or 2 HACs) of each indicated HT1080 subclone 48h after transfection with the indicated fusion proteins; the presence of the HAC was detected by GFP signal. n = 500 cells analyzed per condition. (****P < 0.001; Fisher’s exact test).

Targeting KAP1 to the *α*21-II^LacO/Gal4^ array also resulted in a decrease in HAC-associated CENP-A, although the effect was milder than observed with tethering to the *α*21-I^TetO^ array (Figure 7A). Thus, even though most CENP-A was associated with the *α*21-I^TetO^ region, targeting proteins to the *α*21-II^LacO/Gal4^ region still affected CENP-A levels. This confirms the proximity of the arrays and is consistent with the pattern of histone modifications observed in Figure 5A. The increase in H3K9me3 levels seen after tethering KAP1 to *α*21-II^LacO/Gal4^ array appeared to be more significant than tethering KAP1 to *α*21-I^TetO^. This could be explained by the initial lower level of H3K9me3 on the *α*21-II^LacO/Gal4^ array (Figure 6A, B, C): there might be more unmodified H3K9 that can be converted to H3K9me3 upon the effect of KAP1.

Targeting KAP1 to both arrays simultaneously did not completely suppress kinetochore function as revealed by CENP-A levels, which are partly maintained. In the double tethering, neither CENP-A nor H3K9me3 levels differed greatly from the single tethering, rejecting the hypothesis that the arrays are independent and they would cooperate to establish a state of “super-repression” when both targeted with KAP1 (Figure 7 A, B).

In parallel with measuring the effects of KAP1 tethering on CENP-A and H3K9me3 levels, we also scored the effects of this tethering on centromere function (e.g., HAC segregation in mitosis). Despite differences in levels of correctly or mis-segregating HACs in the initial cell populations, targeting KAP1 to one or both arrays always led to a significant increase in the number of mis-segregating HACs. Interestingly, segregation was significantly impaired even when over-all CENP-A levels were not greatly reduced by KAP1 (Figure 7C). This is probably because the percentage of mis-segregating cells is determined by scoring individual cells in which the level of CENP-A falls below a critical threshold, and it is not determined by the average CENP-A level in the cell population (consistent with Martins et al, unpublished).

Together, these observations lead to the conclusion that CENP-A, H3K4me2 and H3K36me2, which are all necessary for kinetochore maintenance and function, are enriched on the *α*21-I^TetO^ array in the alphoid^2domain^ HAC. Nevertheless, the close proximity and scrambled structure of the two arrays allows the chromatin modifier KAP1 to act simultaneously on both arrays.

## Discussion

We have generated several alphoid^2domain^ HACs by transfecting HT1080 cells with pBAC11.32TW12.32GLII, a HAC-seeding DNA of ∼120 kb. This HAC-seeding DNA contains two distinct *α*-satellite DNA arrays: one rich in binding sites for TetR and CENP-B and one lacking CENP-B boxes but having binding sites for LacI and Gal4. We had expected that the former might form centrochromatin and the latter heterochromatin, but experimental results revealed another outcome.

The new alphoid^2domain^ HAC shows two important differences from previous generations of synthetic HACs (alphoid^tetO^ HAC and alphoid^hybrid^ HAC)^2,4^. First, the efficiency of alphoid^2domain^ HAC formation in HT1080 was higher than that typically seen with other HACs^2,4^. Indeed, it was comparable to results obtained when co-transfecting the HAC-seeding DNA plus CENP-A specifically targeted to the synthetic centromere^4^. It therefore appears that this longer HAC-seeding DNA may be more efficient at promoting stable CENP-A deposition. Second, ChIP-qPCR analysis revealed that CENP-A accumulated preferentially on the CENP-B-containing array of the alphoid^2domain^ HAC. This was not observed with the previous alphoid^hybrid^ HAC, which was formed from a smaller HAC-seeding DNA^4^.

Surprisingly, H3K9me3 was also recruited to the CENP-B-containing array on the alphoid^2domain^ HAC. Previous results have revealed that CENP-B can have a dual role in recruiting centrochromatin or heterochromatin markers depending on the context^10,27^. We speculate that the alphoid^2domain^ HAC shows 3 types of chromatin. Some of the CENP-B-rich arrays form classical centrochromatin^39^ containing CENP-A, H3K4me2 and H3K36me2, but others form H3K9me3-rich heterochromatin, which previous studies have shown to be incompatible with centrochromatin. Thus, CENP-A-containing arrays are likely interspersed with H3K9me3-bearing arrays. Surprisingly, the non-CENP-B array did not form the predicted heterochromatin, but instead appeared to form relatively “open” euchromatin, possibly as a result of selective pressure to avoid silencing the geneticin resistance gene.

Studies of the alphoid^tetO^ HAC revealed that HAC-seeding DNA can undergo dramatic reorganization during HAC establishment^3^. However, the timing and the causes of this phenomenon were unknown. Our Southern blot analysis of various clones of alphoid^2domain^ HAC-bearing HT1080 cells reveals that the HAC-seeding DNA in each clone has undergone a unique pattern of rearrangements, both in the size and in the number of fragments observed after restriction digestion and probing for the arrays present in pBAC11.32TW12.32GLII. This highly cell-specific pattern is acquired by each cell in the first 8 weeks following transfection with the HAC-seeding DNA, apparently before completion of the multimerization/amplification that allows the transfected DNA to surpass the size threshold required for stable segregation in mitosis^23^. The specific pattern of rearrangements is stably inherited by HAC-containing subclones, as shown by Southern blot analysis performed 14 weeks after HAC seeding DNA transfection and in agreement with previous reports of HAC stability during MMCT (Microcell Mediated Cell Transfer)^3,35^. These observations indicate that early during the process of centromere formation, the HAC-seeding DNA encounters a limited series of events that lead to deletions, additions and shuffling of its arrays, but that subsequent to centromere formation the HAC genome is stabilized.

We propose three discrete steps at which modifications on the HAC-seeding DNA possibly occur: in the cytosol, shortly after entry of the HAC-seeding DNA (1st step), in the nucleus during replication (2nd step) and as a consequence of micronucleus formation (3rd step) (Figure 8). Not every HAC-seeding DNA will necessarily undergo all three steps, but we suggest that they can all cooperate to form the rearranged mature HAC.

**Figure 8:**
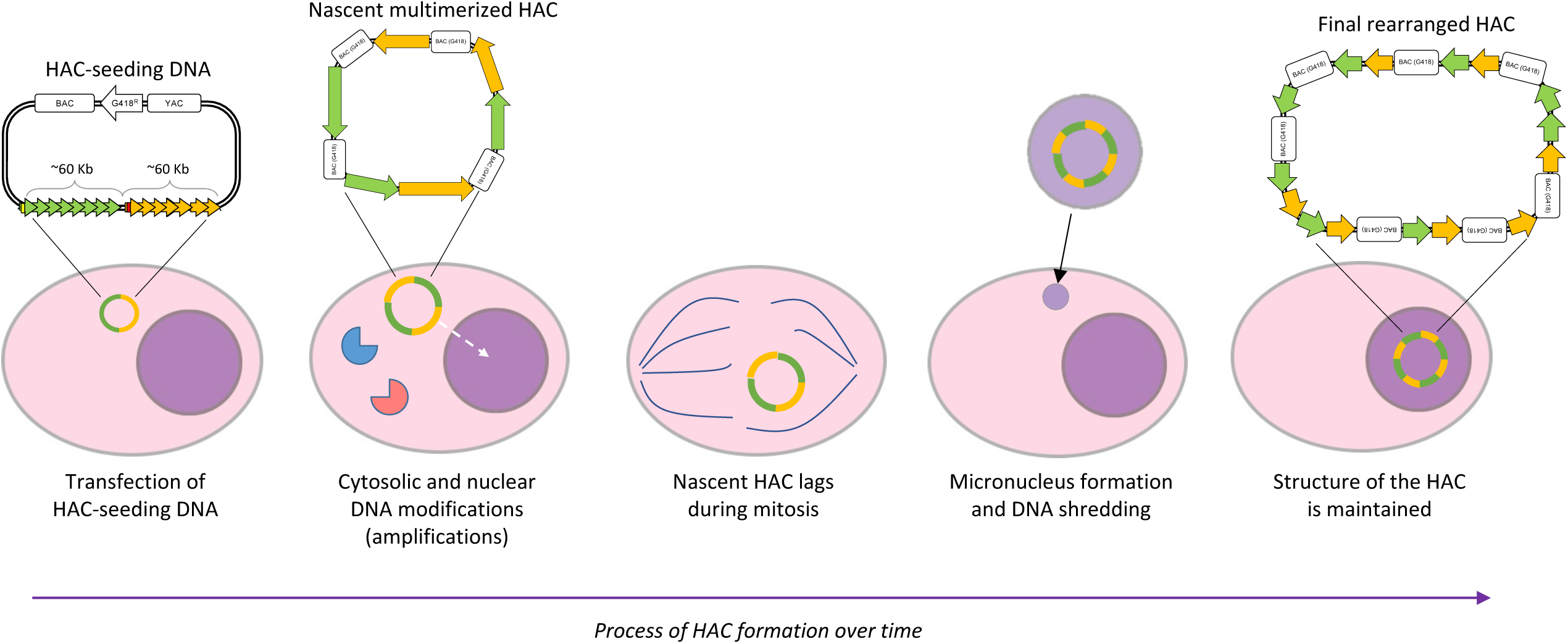
Scheme representing how HAC-seeding DNA rearranges to form the alphoid^2domain^ HAC. HAC-seeding DNA formed by *α*21-I^TetO^ (green) and *α*21-II^LacO/Gal4^ (yellow) arrays is transfected in HT1080 cells; under the effect of proteins of the DNA-sensing pathways (elements in red and blue), the nascent HAC gets initially rearranges and it increases in size due to “slippage” during replication (dashed arrow indicates nascent HAC entering the nucleus). During mitosis, the nascent HAC lags and it gets therefore incorporated into a micronucleus, resulting in additional massive rearrangements which lead to the final structure of the alphoid^2domain^ HAC.

It has been reported extensively in the literature that exogenous DNA is naturally altered upon transfection into cells. Transfected exogenous DNA can undergo mutations, deletions, formation of concatemers or be eventually lost prior to entering the nucleus^40–45^. Most of these modifications happen early after transfection, before the transfected DNA replicates^46^. This is the result of DNA-sensing pathways^47^: for example, the DNA sensor cGAS binds cytosolic DNA and produces the second messenger cGAMP, which binds STING and leads to activation of an inflammatory response in the nucleus^48–50^. Indeed, it was recently reported that cGAS has an affinity for *α*-satellite DNA^51^. Transfected DNA can also undergo double strand breaks (DSBs) triggering a DNA damage repair response^42^ that can involve non-homologous end joining (NHEJ) or homologous recombination (HR).

We suggest that transfected HAC-seeding DNA activates these cytosolic responses and that this results in an initial round of DNA alterations (1st step in our proposed model). This rearranged DNA then enters the nucleus where it undergoes a process of amplification either due to recombination or “slippage” during replication due to its repetitive sequence, leading to the formation of concatemers (2nd step). If, during these early stages the HAC-seeding DNA fails to reach a length sufficient to establish a functional centromere or accurate regulation of sister chromatid cohesion, then at the subsequent mitosis the nascent HAC would fail to segregate properly, likely ending up as a lagging chromosome in anaphase. Such lagging chromosomes typically lead to chromosome bridges and micronucleus formation^32,52,53^. Both outcomes have been associated with chromothripsis, a disruptive event of shredding and shuffling of the DNA, that is associated with cancer development^30–32,54^. During chromothripsis, a region of the genome is cut in tens to hundreds of pieces by an as-yet unknown agent (although rupture of the nuclear envelope in micronuclei has been reported to lead to cGAS accumulation^55^), and the fragments are re-joined randomly by NHEJ, generating a patchwork of DNA fragments^30^.

If the alphoid^2domain^ HAC -seeding DNA undergoes chromothripsis, large numbers of rearrangements could potentially occur in a very short period of time (3rd step). If the rearranged HAC subsequently attains the minimum size for stable mitotic segregation, this could explain the origin of sister HAC-containing cell lines, each with a unique set of rearrangements. This model suggests that the fate of the HAC seeding-DNA may depend on the phase in the cell cycle when transfection happens, as this could result in a longer or shorter exposure of the DNA molecule to cytosol. Furthermore, NHEJ is more active during the G1 phase of the cell cycle, when sister chromatids are not yet formed^56^. The question of the relationship between HAC DNA re-organization and the cell cycle timing of DNA transfection remains an important one for future study.

The results described here have important implications for ongoing efforts to build synthetic human chromosomes by de novo synthesis^57^, as has been done with great success for budding yeast^58–60^. Unlike budding yeast, which has a point centromere^61^, metazoans have regional centromeres that require establishment of a proper epigenetic environment for their function and stability^62–67^. Our data suggest that this process of centromere formation is frequently associated with DNA rearrangements. It would be extremely unfortunate if human chromosomes synthesized at great cost and effort were to become scrambled in an uncontrollable fashion upon their introduction to human host cells during the process of centromere establishment.

Importantly, once centromere function is established, the associated DNA arrays appear to be much more stable. In fact, although some minor mutations were detected in some clones when the ∼90 kb *BRCA1* gene was inserted into an established HAC vector (possibly the result of step 1), there was no detected chromothripsis^12^. Therefore, alternative strategies for building synthetic chromosomes – e.g. assembly of synthetic human chromosomes by building upon available HACs with a multi-integrase site adjacent to the tetO-array – may avoid these complex DNA rearrangements^68^.

It will be important in future studies to use the alphoid^2domain^ HAC system to establish suitable conditions for conservation of the organization of chromosome-sized DNA molecules introduced into human cells during centromere establishment. Future studies with HACs will allow us to determine whether transfected HAC-seeding DNA does undergo chromothripsis, and if so, how to minimize this. The HAC system will also be useful for studies to optimize procedures to increase the efficiency of centromere activation and establishment of properly regulated cohesion on exogenous DNA. Only when these technical issues have been resolved will it be possible to form pre-determined artificial and synthetic chromosomes in human cells.

## Methods

### Construction of BAC carrying *α*21-I^TetO^ and *α*21-II^LacO/Gal4^ arrays by tandem ligation amplification

The BAC clone pBAC11.32TW12.32GLII, carrying 32 copies of the *α*21-II^LacO/Gal4^ 12-mer following 32 copies of the *α*21-I^TetO^ 11-mer was constructed in three steps describing below.

1. Construction of *α*21-II^LacO/Gal4^ alphoid 12-mer and insertion into pBluescript vector: *α*21-II^LacO/Gal4^ alphoid 12-mer has been designed base on alphoid type II DNA of chromosome #21 and it has been synthesized by GENEART. The SpeI and NheI sites are located respectively at left and right ends of the *α*21-II^LacO/Gal4^ 12-mer to be inserted into pBlurescript vector. The vector and the 12-mer were joined using the homologous recombination-based method (GENEART Seamless Cloning and Assembly Kit, ThermoFisher Scientific). The resulting plasmid carries one copy of *α*21-II^LacO/Gal4^ 12-mer accompanying unique NheI and unique SpeI site at the ends.
2. Extension of the *α*21-II^LacO/Gal4^ 12-mer insert in the plasmid vector by repeating the tandem ligation. To extend length of the alphoid insert, the tandem ligation was repeated until the plasmid harbored 8 copies of the 12-mer using SpeI, NheI and ScaI restriction enzymes (NEB)^29,69^. Therefore, the band of the highest molecular weight (16.6kb for 8 copies) was excised after PFGE and cloned into the BAC vector. The *α*21-I^TetO^ 11-mer was designed based on the sequence of type I alphoid 11-mer of chromosome #21 centromere^21^.
3. Extension of the *α*21-II^LacO/Gal4^ 12-mer insert in the BAC vector. Starting from the BAC clone carrying 8 copies of the *α*21-II^LacO/Gal4^ 12-mer, the tandem ligation was repeated until the 12-mer insert reached 32 copies using SpeI, NheI and KasI restriction enzymes (NEB). Finally, 32 copies of *α*21-I^TetO^ 11-mer has been cut out from the BAC vector and ligated into the same vector of *α*21-II^LacO/Gal4^ 12-mer to obtain the final product, pBAC11.32TW12.32GLII.

After each cloning step, the forming arrays were digested with BamHI and NotI restriction enzymes (NEB) and analysed on 1% agarose gel electrophoresis using 100 bp DNA Ladder, Quick-Load 1 kb Extend DNA ladder or Low Range PFG Marker (NEB).

### Quantitative PCR (qPCR) to detect BAC copy number

Cells from HAC-containing clones have been harvested and genomic DNA has been collected using Maxwell DNA purification kit (Promega). qPCR analysis have been performed using SYBR Green Master Mix (Roche) and the following primers: N11F5: 5’-GGGATCACTAGCAATAAAAGGTAGAC-3’ and N11R6: 5’-TCCTTCTGTCTCGTTTTTATGGC-3’ for the BAC synthetic DNA; 11-10R: 5’-AGG GAA TGT CTT CCC ATA AAA ACT-3’and mCbox-4: 5’-GTC TAC CTT TTA TTT GAA TTC CCG-3’ for the alphoid chr21 array as control. As a standard, DNA from a previously characterized HAC-containing cell line (H21) with a known number of BAC copies (n = 125) has been diluted with serial dilutions and amplified with the same primers.

### Cell culture, transfection, HAC formation and subcloning

Human HT1080 cells were cultured in DMEM supplemented with 10% FBS (Labtech) plus 100 U/ml penicillin G and 100 μg/mL of streptomycin sulfate (Invitrogen). Cells were grown at 37 °C in 5%CO_2_ in a humidified atmosphere. Transfection of pBAC11.32TW12.32GLII DNA was performed using Viafect (Promega) following the manufacturer’s instructions. For transfections of cells growing in 6 wells-plates, transfection complexes containing 10 μL of Viafect reagent and 1 μg of plasmid DNA were prepared in 200 μL of OptiMEM (Invitrogen). After 5 min of incubation at room temperature, 200 μL of transfection complexes was added dropwise in 2 mL of media. After 6 h, the media was changed to the wells and transfected cells were selected adding 400 μg/mL of geneticin (Thermo Fisher) and grown for 2–3 weeks until separate resistant colonies were present. Resistant colonies were isolated manually and moved into 24 wells-plates. Isolated clones were expanded in the presence of 400 μg/mL of geneticin. For targeting experiments with TerR-KAP1 and LacI-KAP1, cells have been transfected using Xtremegene-9 (Roche) according to manifacturer’s instructions. For transfections in 12 wells-plates, transfection complexes containing 3 μL of Xtremegene-9 reagent and 500 ng of plasmid DNA were prepared in 100 μL of OptiMEM (Invitrogen). After 20 min of incubation at room temperature, 100 μL of transfection complexes was added dropwise in 1 mL of media. For co-transfection, 500 ng of each plasmid has been transfected in the same reaction.

### Southern blot hybridization analysis

Southern blot hybridization was performed as described previously^35^ with minor changes. Genomic DNA was prepared in agarose plugs (Low Melt Agarose, Biorad), 0.5×10^6^ cells per plug: plugs have been treated with Proteinase K (CHEF genomic DNA plug kit, Biorad) and restriction-digested by BamHI (NEB) overnight in the buffer recommended by the manufacturer. The digested DNA was CHEF (CHEF Mapper, Bio-Rad) separated (autoprogram, 5–250 kb range, 16 h transfer), transferred to membrane (Amersham Hybond-N+), and blot-hybridized with an 82 bp probe for TetO and a 74 bp probe for LacO and Gal4 containing P-32 CTP. Radioactive labelled DNA sequences for the probes were synthesized by PCR using the primers and synthetic DNA fragment as a template (tetO_south_21_M:5’TTTGTGGAAGTGGACATTTACTAGCAGCAGAGCTCTCCCT ATCAGTGATAGAGACTAGCCCATAA AATAGACAGAAGCATT-3′, tetO_south_21_F1:5′-TTTGTGGAAGTGGACATTTC-3′, tetO_south_21_R1:5′-AATGCTTCTGTCTATTTTTA-3′;lacO_south3_M:5′TGTGGAAGTGGACATTTCGACCACATGTGGAATTGTGAGCGGATAACAATTTGTGGCCCATAAAATAGACAGA-3′, lacO_south3_F1: 5′-TGTGGAAGT GGACATTTCGA-3′, lacO_south3_R1: 5′-TCTGTCTATTTTTATGGGCC-3′; gal4_south1_M: 5′-AATGGACATTTCGACGGAGGACAGTCCTCCGTCGACGGAGG ACAGTCCTCCGCATAAAATCTA-3′,Gal4_south1_F1: 5′-AATGGACATTTCGACG-3′, gal4_south1_R1: 5′-TAGATTTTATGCGGAG-3′).

The membrane was incubated for 2 h at 65 °C for prehybridization in Church’s buffer (0.5 M Na-phosphate buffer containing 7% SDS and 100 μg/mL of unlabeled salmon sperm carrier DNA). The labelled probe was heat denatured in a boiling water for 5 min, cooled, added to the hybridization Church’s buffer and allowed to hybridize for 48 h at 65 °C. Blots were washed once in 2x SSC (300 mM NaCl, 30 mM sodium citrate, pH 7.0)/0.05% SDS for 20 min at 30 °C, once in 2x SSC/0.05% SDS for 10 min at 65 °C and then three times in 2x SSC/ 0.05% SDS for 5 min at 65 °C. Blots were exposed to X-ray film 2-48hrs. at -80 °C.

### Expression and purification of recombinant TetR/LacI-eYFP

TetR and LacI were cloned as C-terminally His-tagged proteins in a pET23a vector and proteins were purified following a previously described procedure^9^. Briefly, the vectors were transformed in *E. coli* BL21 Gold cells and colonies grown at 37°C until OD_600_ 1 in Super Broth containing ampicillin. The cultures were then induced with 0.35 mM IPTG overnight at 18°C and cell pellets were lysed in a buffer containing 20 mM Tris HCl pH 7.5, 500 mM NaCl, 35 mM imidazole and 2 mM 2-mercaptoethanol. Proteins were affinity-purified using a Ni-NTA column (GE Healthcare), washed with high salt buffer (20 mM Tris HCl pH 7.5, 1000 mM NaCl, 50 mM KCl, 10 mM MgCl_2_, 2 mM ATP, 35 mM imidazole and 2 mM 2-mercaptoethanol) and eluted with 20 mM Tris HCl pH 7.5, 150 mM NaCl, 400 mM imidazole and 2 mM 2-mercaptoethanol. The pure eluted fractions were pooled and dialysed overnight against storage buffer (20 mM Tris HCl pH 7.5, 150 mM NaCl, 5% glycerol and 2 mM 2-mercaptoethanol). Sample quality was analysed by 15% SDS-PAGE stained with Coomassie Blue. The final protein concentrations that were used for the immuno-fiber-FISH were 1.2 and 1.7 mg/ml for TetR-eYFP and LacI-eYFP, respectively.

### Fluorescent in situ hybridization (FISH) and Immuno-fiber-FISH

Metaphase chromosomes from HT1080 were obtained following a standard protocol: 3 h before harvesting, cells were treated with Colcemid (Invitrogen) at a final concentration of 0.1 μg/mL. Collected cells were resuspended in warm hypotonic solution (75mM KCl) for 20 min at 37 °C and fixed in methanol:acetic acid (3:1). Slides were kept at –20 °C until they were processed for FISH. To obtain stretched chromatin fibers, 2×10^6^ cells were centrifuged, and the pellets were washed in 1xPBS. 10 µl -drops have been placed on slides and let dry. Once the slides were mounted on the Shandon Sequenza cover plates (Thermo Scientific), DNA fibers were released applying a lysis solution (700mM NaOH in ethanol) and fixed in methanol. Slides have been kept in PBS at 4 °C until stained.

For oligo-FISH staining, oligonucleotides recognizing the tetO sequence (5′-ACTAGCAGCAGAGCTCTCCCTATCAGTGATAGAGACTAG-3′) labelled with Digoxigenin, and oligonucleotides recognizing both lacO (5′-CATGTGGAATTGTGAGCGGATAACAATTTGTGG-3′) and Gal4 (5′-TCGACGGAGGACAGTCCTCCG-3′) sequences labelled with Biotin were purchased (Sigma). Oligonucleotides were mixed at 100 ng/μL and resuspended in hybridization buffer (50% formamide, 10% dextran sulfate, 2x SSC (300 mM NaCl, 30 mM sodium citrate, pH 7.0)) and 50 μg/mL of salmon sperm DNA (Sigma). FISH was carried out following standard procedures. Slides were denatured in 70% formamide/2x SSC at 70 °C for 45 sec and hybridized in a humid chamber at 37 °C for 2 h. Slides were then washed in 20%formamide/2x SSC for 5 min and in 2x SSC/0.1%Tween-20 for 5 min at 37 °C. Oligonucleotide probes were detected with rhodamine-conjugated antidigoxigenin (Roche) and fluorescein-conjugated streptavidin (Vector Laboratories) incubated for 30 min at 37 °C. Slides were mounted with Vectashield (Vector Laboratories) containing 4′,6-diamidino-2-phenylindole (DAPI) for chromosome counterstaining. For immune-fiber-FISH, slides have been blocked with 1% BSA/1xPBS/0.1% Triton x-100 and then incubated with TetR-GFP or LacI-GFP proteins purified from *E*.*Coli* in blocking buffer overnight at 4 °C, together with *α*-CENP-A and *α*-H3K9me3 antibodies as described in immunofluorescence (IF) protocol. Slides have been washed, incubated with secondary antibodies and sealed as immunofluorescence (IF) staining standard protocol.

### Analysis of the HAC stability

HAC-containing subclones have been thawed and maintained in culture with 400 μg/mL of geneticin (Thermo Fisher) for 7 days. At day 0, metaphase chromosomes have been spread on slides and labelled for FISH as described. At day 0, cell cultures have been split into two batches: one batch has been kept in culture with 400 μg/mL of geneticin (Thermo Fisher) for 30 days, while the other has been kept in culture with simple DMEM/10% FBS/1% PenStrepto. At day 30, metaphase chromosomes from each batch have been spread on slides and labelled for FISH as described. Metaphases at day 0 and day 30 have been scored for the presence of 0,1, 2 or >2 HACs. The daily loss rate of the HAC (R) was calculated using the formula Nn= N0 × (1 – R)^n^, where N0 is the number of metaphase chromosome spreads showing a HAC in the cells cultured under selection and Nn is the number of HAC-containing metaphase chromosome spreads after n days of culture in the absence of selection.

### Indirect immunofluorescence (IF) staining and microscopy analysis

Indirect immunofluorescence (IF) staining of cells fixed in 3.7% formaldehyde/1x PBS was performed at 37°C for 10 minutes following standard procedures. The following antibodies were used: mouse anti-CENP-A (clone A1, 1:500,^36^) and rabbit anti-H3K9me3 (abcam 8898; 1:200). Microscope images were acquired on a DeltaVision Core system (Applied Precision) using an Olympus IX-71 inverted microscope stand with an Olympus UPlanSApo 100x oil immersion objective (numerical aperture (NA) 1.4) and an LED light source. Camera (Photometrics Cool Snap HQ), shutter, and stage were controlled through SoftWorx (Applied Precision). Z-series were collected with a spacing of 0.2 μm, and image stacks were subsequently deconvolved in SoftWorx. CENP-A and H3K9me3 signal quantification at the HAC was performed using ImageJ. HAC was visualized in the green channel thanks to the tethering of the corresponding fluorescent protein. For CENP-A signal quantification, a custom-made macro in ImageJ (National Institutes of Health, Bethesda, MD) was used^70^.

### Chromatin Immunoprecipitation and quantitative PCR (ChIP-qPCR)

Exponentially growing cells were harvested with TrypLE Express (Gibco) and resuspended in D-PBS (Gibco) up to a concentration of 1x 10^6^ cells/ml. They were cross-linked with 1% formaldehyde solution (Sigma) for 5 min at room temperature, followed by quenching with 2.5 M glycine for 5 min at room temperature. Cells at a concentration of 5x 10^6^ cells/ml were lysed in lysis buffer (10 mM Tris pH 8.0; 10 mM NaCl; 0.5% NP-40) containing protease inhibitors (1 μg/mL of CLAP; 0.5 μg/mL of Aprotinin; 1 mM PMSF) for 10 min on ice. Cells pellets have been snap-freezed in liquid N and stored in - 80 °C until processed. Nuclei were resuspended in lysis buffer with protease inhibitors (50 mM Tris pH 8.0; 2 mM EDTA; 0.2% SDS; 134 mM NaCl; 0.88% Triton X-100; 0.088% Na-deoxycholate) and incubated with 400U/ml MNase for 30 min at 20 °C. Chromatin was sheared by sonication in a Bioruptor sonicator (Diagenode) for 9 cycles (30s ON/30S OFF) at high setting and 4 °C in dilution buffer 1 enriched with. The collected supernatants after sonication were diluted with 300 μL of dilution buffer 1, 500 μL of dilution buffer 2 (50 mM Tris pH 8.0; 167 mM NaCl; 1.1% Triton X-100; 0.11% Na-deoxycholate), and 500 μL of RIPA buffer containing 150 μL of NaCl (RIPA-150) and protease inhibitors. Antimouse IgG Dynabeads (Invitrogen) were conjugated with the relevant antibodies for 6 h with RIPA-150/0.5% BSA at 4 °C and washed twice with RIPA-150/0.5% BSA. 500 μL of collected sheared chromatin was incubated with the beads at 4 °C overnight. Beads were afterward washed twice with RIPA-150 and RIPA buffer containing 500 mM NaCl (RIPA-500) and a final wash with TE pH 8.0. Antibody/chromatin complexes were de-cross-linked with 10% Chelex-100 resin (BioRad) in water at 93 °C and incubated with RNase A and Proteinase K. DNA was subsequently recovered using magnetic rack. ChIPed DNA was subjected to RT-PCR using a SYBR Green Master Mix (Roche) using the following oligonucleotides: tetO-Fw (5′-CCACTCCCTATCAGTGATAGAGAA-3′), tetO-Rv (5′-TCGACTTCTGTTTAGTTCTGTGCG-3′) for the α21-I-tetO domain of the hybrid HAC, lacOgal4-Fw (5′-TATGGTGTCGACGGAGGACA-3′), and lacOgal4-Rv (5′-CCGCTCACAATTCCACATGTG-3′) for the α21-II-lacOgal4 domain of the hybrid HAC, chr17-Fw (5′-TTGTGGTTTGTGGTGGAAAA-3′) and chr17-Rv (5′-CTCAAAGCGCTCCAAATCTC-3′) for the alphoid chr17 array, sat2-Fw (5′ TCGCATAGAATCGAATGGAA-3′) and sat2-Rv (5′-GCATTCGAGTCCGTGGA-3′) for the pericentromeric alphoid chr1.

## Author Information

AM is currently at Telethon Institute of Genetics and Medicine, Pozzuoli, Italy

## Author Contributions

E.P. and W.C.E. designed the experiments; E.P. performed the experiments, with the help of A.M. and C.R.; M.L. performed Southern blot experiments; K.O. constructed the HAC seeding DNA; M.A.A. expressed bacteria purified proteins; M.L., H.M., V.L., N.K. and W.C.E. contributed with critical discussions and new experimental tools; E.P. analyzed the data; E.P. and W.C.E. wrote the paper.

## Acknowledgements

This work was funded by the UK Research Councils’ Synthetic Biology for Growth programme [core grant BB/M018040/1 to the BBSRC/EPSRC/MRC Synthetic Biological Research Centre] (E.P). Additional funding of the Earnshaw lab is provided by Wellcome, of which W.C.E. is a Principal Research Fellow [grant number 073915 Imaging was performed in Centre Optical Instrumentation Laboratory (COIL), which is supported by a Core Grant (203149)] to the Wellcome Centre for Cell Biology at the University of Edinburgh. Additional experiments were supported by MEXT KAKENHI [grant numbers 16H04747, 16H01414 and 18H04721] and the Kazusa DNA Research Institute Foundation (H.M.) and by the Intramural Research Program of the NIH, National Cancer Institute, Center for Cancer Research, USA(V.L.). MAA and AAJ are supported by the Wellcome Trust through a Wellcome Senior Research Fellowship to AAJ (202811). We also thank Fernanda Cisneros Soberanis for her feedback on the manuscript.

## Graphical abstract

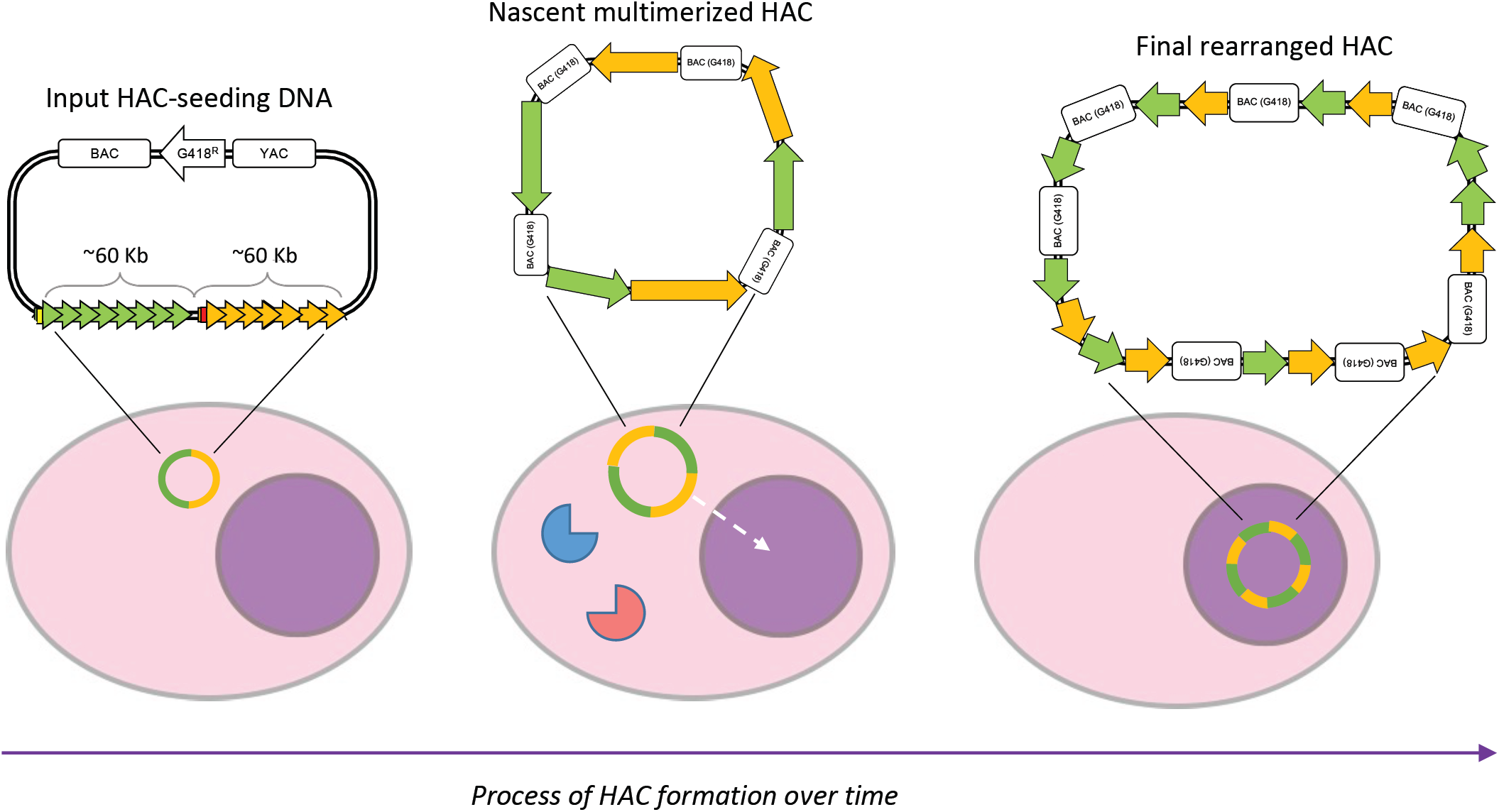

## References

(1) Harrington, J. J.; Van Bokkelen, G.; Mays, R. W.; Gustashaw, K.; Willard, H. F.Formation of de Novo Centromeres and Construction of First-Generation Human Artificial Microchromosomes. Nat. Genet. 1997. https://doi.org/10.1038/ng0497-345.

(2) Nakano, M.; Cardinale, S.; Noskov, V. N.; Gassmann, R.; Vagnarelli, P.; Kandels-Lewis, S.; Larionov, V.; Earnshaw, W. C.; Masumoto, H. Inactivation of a Human Kinetochore by Specific Targeting of Chromatin Modifiers. Dev. Cell 2008. https://doi.org/10.1016/j.devcel.2008.02.001.

(3) Kouprina, N.; Samoshkin, A.; Erliandri, I.; Nakano, M.; Lee, H. S.; Fu, H.; Iida, Y.; Aladjem, M.; Oshimura, M.; Masumoto, H.; Earnshaw, W. C.; Larionov, V. Organization of Synthetic Alphoid DNA Array in Human Artificial Chromosome (HAC) with a Conditional Centromere. ACS Synth. Biol. 2012. https://doi.org/10.1021/sb3000436.

(4) Pesenti, E.; Kouprina, N.; Liskovykh, M.; Aurich-Costa, J.; Larionov, V.; Masumoto, H.; Earnshaw, W. C.; Molina, O. Generation of a Synthetic Human Chromosome with Two Centromeric Domains for Advanced Epigenetic Engineering Studies. ACS Synth. Biol. 2018. https://doi.org/10.1021/acssynbio.8b00018.

(5) Lee, N. C. O.; Kim, J. H.; Petrov, N. S.; Lee, H. S.; Masumoto, H.; Earnshaw, W. C.; Larionov, V.; Kouprina, N. Method to Assemble Genomic DNA Fragments or Genes on Human Artificial Chromosome with Regulated Kinetochore Using a Multi-Integrase System. ACS Synth. Biol. 2018. https://doi.org/10.1021/acssynbio.7b00209.

(6) Cardinale, S.; Bergmann, J. H.; Kelly, D.; Nakano, M.; Valdivia, M. M.; Kimura, H.; Masumoto, H.; Larionov, V.; Earnshaw, W. C.Hierarchical Inactivation of a Synthetic Human Kinetochore by a Chromatin Modifier. Mol. Biol. Cell 2009, 20(19), 4194–4204. https://doi.org/10.1091/mbc.E09-06-0489.

(7) Bergmann, J. H.; Rodríguez, M. G.; Martins, N. M. C.; Kimura, H.; Kelly, D. A.; Masumoto, H.; Larionov, V.; Jansen, L. E. T.; Earnshaw, W. C.Epigenetic Engineering Shows H3K4me2 Is Required for HJURP Targeting and CENP-A Assembly on a Synthetic Human Kinetochore. EMBO J. 2011. https://doi.org/10.1038/emboj.2010.329.

(8) Bergmann, J. H.; Jakubsche, J. N.; Martins, N. M.; Kagansky, A.; Nakano, M.; Kimura, H.; Kelly, D. A.; Turner, B. M.; Masumoto, H.; Larionov, V.; Earnshaw, W. C.Epigenetic Engineering: Histone H3K9 Acetylation Is Compatible with Kinetochore Structure and Function. J. Cell Sci. 2012. https://doi.org/10.1242/jcs.090639.

(9) Molina, O.; Vargiu, G.; Abad, M. A.; Zhiteneva, A.; Jeyaprakash, A. A.; Masumoto, H.; Kouprina, N.; Larionov, V.; Earnshaw, W. C.Epigenetic Engineering Reveals a Balance between Histone Modifications and Transcription in Kinetochore Maintenance. Nat. Commun. 2016. https://doi.org/10.1038/ncomms13334.

(10) Ohzeki, J. I.; Bergmann, J. H.; Kouprina, N.; Noskov, V. N.; Nakano, M.; Kimura, H.; Earnshaw, W. C.; Larionov, V.; Masumoto, H. Breaking the HAC Barrier: Histone H3K9 Acetyl/Methyl Balance Regulates CENP-A Assembly. EMBO J. 2012. https://doi.org/10.1038/emboj.2012.82.

(11) Kim, J. H.; Kononenko, A.; Erliandri, I.; Kim, T. A.; Nakano, M.; Iida, Y.; Barrett, J. C.; Oshimura, M.; Masumoto, H.; Earnshaw, W. C.; Larionov, V.; Kouprina, N. Human Artificial Chromosome (HAC) Vector with a Conditional Centromere for Correction of Genetic Deficiencies in Human Cells. Proc. Natl. Acad. Sci. U. S. A. 2011. https://doi.org/10.1073/pnas.1114483108.

(12) Kononenko, A. V.; Bansal, R.; Lee, N. C. O.; Grimes, B. R.; Masumoto, H.; Earnshaw, W. C.; Larionov, V.; Kouprina, N. A Portable BRCA1-HAC (Human Artificial Chromosome) Module for Analysis of BRCA1 Tumor Suppressor Function. Nucleic Acids Res. 2014. https://doi.org/10.1093/nar/gku870.

(13) Kouprina, N.; Tomilin, A. N.; Masumoto, H.; Earnshaw, W. C.; Larionov, V. Human Artificial Chromosome-Based Gene Delivery Vectors for Biomedicine and Biotechnology. Expert Opinion on Drug Delivery. 2014. https://doi.org/10.1517/17425247.2014.882314.

(14) Kononenko, A. V.; Lee, N. C. O.; Liskovykh, M.; Masumoto, H.; Earnshaw, W. C.; Larionov, V.; Kouprina, N. Generation of a Conditionally Self-Eliminating HAC Gene Delivery Vector through Incorporation of a TTAVP64 Expression Cassette. Nucleic Acids Res. 2015. https://doi.org/10.1093/nar/gkv124.

(15) Liskovykh, M.; Ponomartsev, S.; Popova, E.; Bader, M.; Kouprina, N.; Larionov, V.; Alenina, N.; Tomilin, A. Stable Maintenance of de Novo Assembled Human Artificial Chromosomes in Embryonic Stem Cells and Their Differentiated Progeny in Mice. Cell Cycle 2015. https://doi.org/10.1080/15384101.2015.1014151.

(16) Ponomartsev, S. V.; Sinenko, S. A.; Skvortsova, E. V.; Liskovykh, M. A.; Voropaev, I. N.; Savina, M. M.; Kuzmin, A. A.; Kuzmina, E. Y.; Kondrashkina, A. M.; Larionov, V.; Kouprina, N.; Tomilin, A. N.Human AlphoidtetO Artificial Chromosome as a Gene Therapy Vector for the Developing Hemophilia A Model in Mice. Cells 2020. https://doi.org/10.3390/cells9040879.

(17) Lee, H. S.; Lee, N. C. O.; Kouprina, N.; Kim, J. H.; Kagansky, A.; Bates, S.; Trepel, J. B.; Pommier, Y.; Sackett, D.; Larionov, V. Effects of Anticancer Drugs on Chromosome Instability and New Clinical Implications for Tumor-Suppressing Therapies. Cancer Res. 2016. https://doi.org/10.1158/0008-5472.CAN-15-1617.

(18) Lee, H. S.; Carmena, M.; Liskovykh, M.; Peat, E.; Kim, J. H.; Oshimura, M.; Masumoto, H.; Teulade-Fichou, M. P.; Pommier, Y.; Earnshaw, W. C.; Larionov, V.; Kouprina, N. Systematic Analysis of Compounds Specifically Targeting Telomeres and Telomerase for Clinical Implications in Cancer Therapy. Cancer Res. 2018. https://doi.org/10.1158/0008-5472.CAN-18-0894.

(19) Liskovykh, M.; Goncharov, N. V.; Petrov, N.; Aksenova, V.; Pegoraro, G.; Ozbun, L. L.; Reinhold, W. C.; Varma, S.; Dasso, M.; Kumeiko, V.; Masumoto, H.; Earnshaw, W. C.; Larionov, V.; Kouprina, N. A Novel Assay to Screen SiRNA Libraries Identifies Protein Kinases Required for Chromosome Transmission. Genome Res. 2019. https://doi.org/10.1101/gr.254276.119.

(20) Ikeno, M.; Suzuki, N. Construction and Use of a Bottom-up HAC Vector for Transgene Expression. Methods Mol. Biol. 2011. https://doi.org/10.1007/978-1-61779-099-7_7.

(21) Ikeno, M.; Grimes, B.; Okazaki, T.; Nakano, M.; Saitoh, K.; Hoshino, H.; McGill, N. I.; Cooke, H.; Masumoto, H. Construction of YAC-Based Mammalian Artificial Chromosomes. Nat. Biotechnol. 1998. https://doi.org/10.1038/nbt0598-431.

(22) Logsdon, G. A.; Gambogi, C. W.; Liskovykh, M. A.; Barrey, E. J.; Larionov, V.; Miga, K. H.; Heun, P.; Black, B. E.Human Artificial Chromosomes That Bypass Centromeric DNA. Cell 2019. https://doi.org/10.1016/j.cell.2019.06.006.

(23) Farr, C. J.; Bayne, R. A.; Kipling, D.; Mills, W.; Critcher, R.; Cooke, H. J.Generation of a Human X-Derived Minichromosome Using Telomere-Associated Chromosome Fragmentation. EMBO J. 1995. https://doi.org/10.1002/j.1460-2075.1995.tb00228.x.

(24) Earnshaw, W. C.; Sullivan, K. F.; Machlin, P. S.; Cooke, C. A.; Kaiser, D. A.; Pollard, T. D.; Rothfield, N. F.; Cleveland, D. W.Molecular Cloning of CDNA for CENP-B, the Major Human Centromere Autoantigen. J. Cell Biol. 1987. https://doi.org/10.1083/jcb.104.4.817.

(25) Masumoto, H.; Masukata, H.; Muro, Y.; Nozaki, N.; Okazaki, T. A Human Centromere Antigen (CENP-B) Interacts with a Short Specific Sequence in Alphoid DNA, a Human Centromeric Satellite. J. Cell Biol. 1989. https://doi.org/10.1083/jcb.109.5.1963.

(26) Earnshaw, W. C.; Rothfield, N. Identification of a Family of Human Centromere Proteins Using Autoimmune Sera from Patients with Scleroderma. Chromosoma 1985. https://doi.org/10.1007/BF00328227.

(27) Okada, T.; Ohzeki, J. ichirou; Nakano, M.; Yoda, K.; Brinkley, W. R.; Larionov, V.; Masumoto, H. CENP-B Controls Centromere Formation Depending on the Chromatin Context. Cell 2007. https://doi.org/10.1016/j.cell.2007.10.045.

(28) Masumoto, H.; Nakano, M.; Ohzeki, J. I.The Role of CENP-B and α-Satellite DNA: De Novo Assembly and Epigenetic Maintenance of Human Centromeres. Chromosom. Res. 2004. https://doi.org/10.1023/B:CHRO.0000036593.72788.99.

(29) Ohzeki, J. ichirou; Nakano, M.; Okada, T.; Masumoto, H. CENP-B Box Is Required for de Novo Centromere Chromatin Assembly on Human Alphoid DNA. J. Cell Biol. 2002. https://doi.org/10.1083/jcb.200207112.

(30) Stephens, P. J.; Greenman, C. D.; Fu, B.; Yang, F.; Bignell, G. R.; Mudie, L. J.; Pleasance, E. D.; Lau, K. W.; Beare, D.; Stebbings, L. A.; McLaren, S.; Lin, M. L.; McBride, D. J.; Varela, I.; Nik-Zainal, S.; Leroy, C.; Jia, M.; Menzies, A.; Butler, A. P.; Teague, J. W.; Quail, M. A.; Burton, J.; Swerdlow, H.; Carter, N. P.; Morsberger, L. A.; Iacobuzio-Donahue, C.; Follows, G. A.; Green, A. R.; Flanagan, A. M.; Stratton, M. R.; Futreal, P. A.; Campbell, P. J.Massive Genomic Rearrangement Acquired in a Single Catastrophic Event during Cancer Development. Cell 2011. https://doi.org/10.1016/j.cell.2010.11.055.

(31) Liu, P.; Erez, A.; Nagamani, S. C. S.; Dhar, S. U.; Kołodziejska, K. E.; Dharmadhikari, A. V.; Cooper, M. L.; Wiszniewska, J.; Zhang, F.; Withers, M. A.; Bacino, C. A.; Campos-Acevedo, L. D.; Delgado, M. R.; Freedenberg, D.; Garnica, A.; Grebe, T. A.; Hernández-Almaguer, D.; Immken, L.; Lalani, S. R.; McLean, S. D.; Northrup, H.; Scaglia, F.; Strathearn, L.; Trapane, P.; Kang, S. H. L.; Patel, A.; Cheung, S. W.; Hastings, P. J.; Stankiewicz, P.; Lupski, J. R.; Bi, W. Chromosome Catastrophes Involve Replication Mechanisms Generating Complex Genomic Rearrangements. Cell 2011. https://doi.org/10.1016/j.cell.2011.07.042.

(32) Crasta, K.; Ganem, N. J.; Dagher, R.; Lantermann, A. B.; Ivanova, E. V.; Pan, Y.; Nezi, L.; Protopopov, A.; Chowdhury, D.; Pellman, D. DNA Breaks and Chromosome Pulverization from Errors in Mitosis. Nature. 2012. https://doi.org/10.1038/nature10802.

(33) Ebersole, T.; Okamoto, Y.; Noskov, V. N.; Kouprina, N.; Kim, J. H.; Leem, S. H.; Barrett, J. C.; Masumoto, H.; Larionov, V. Rapid Generation of Long Synthetic Tandem Repeats and Its Application for Analysis in Human Artificial Chromosome Formation. Nucleic Acids Res. 2005. https://doi.org/10.1093/nar/gni129.

(34) Ohzeki, J. ichirou; Larionov, V.; Earnshaw, W. C.; Masumoto, H. Genetic and Epigenetic Regulation of Centromeres: A Look at HAC Formation. Chromosom. Res. 2015. https://doi.org/10.1007/s10577-015-9470-z.

(35) Liskovykh, M.; Lee, N. C.; Larionov, V.; Kouprina, N. Moving toward a Higher Efficiency of Microcell-Mediated Chromosome Transfer. Mol. Ther. - Methods Clin. Dev. 2016. https://doi.org/10.1038/mtm.2016.43.

(36) Kimura, H.; Hayashi-Takanaka, Y.; Goto, Y.; Takizawa, N.; Nozaki, N. The Organization of Histone H3 Modifications as Revealed by a Panel of Specific Monoclonal Antibodies. Cell Struct. Funct. 2008. https://doi.org/10.1247/csf.07035.

(37) Schultz, D. C.; Friedman, J. R.; Rauscher, F. J.Targeting Histone Deacetylase Complexes via KRAB-Zinc Finger Proteins: The PHD and Bromodomains of KAP-1 Form a Cooperative Unit That Recruits a Novel Isoform of the Mi-2α Subunit of NuRD. Genes Dev. 2001. https://doi.org/10.1101/gad.869501.

(38) Cardinale, S.; Bergmann, J. H.; Kelly, D.; Nakano, M.; Valdivia, M. M.; Kimura, H.; Masumoto, H.; Larionov, V.; Earnshaw, W. C.Hierarchical Inactivation of a Synthetic Human Kinetochore by a Chromatin Modifier. Mol. Biol. Cell 2009. https://doi.org/10.1091/mbc.E09-06-0489.

(39) Sullivan, B. A.; Karpen, G. H.Centromeric Chromatin Exhibits a Histone Modification Pattern That Is Distinct from Both Euchromatin and Heterochromatin. Nat. Struct. Mol. Biol. 2004. https://doi.org/10.1038/nsmb845.

(40) Wake, C. T.; Gudewicz, T.; Porter, T.; White, A.; Wilson, J. H.How Damaged Is the Biologically Active Subpopulation of Transfected DNA? Mol. Cell. Biol. 1984. https://doi.org/10.1128/mcb.4.3.387.

(41) Würtele, H.; Little, K. C. E.; Chartrand, P. Illegitimate DNA Integration in Mammalian Cells. Gene Therapy. 2003. https://doi.org/10.1038/sj.gt.3302074.

(42) Dellaire, G.; Yan, J.; Little, K. C. E.; Drouin, R.; Chartrand, P. Evidence That Extrachromosomal Double-Strand Break Repair Can Be Coupled to the Repair of Chromosomal Double-Strand Breaks in Mammalian Cells. Chromosoma 2002. https://doi.org/10.1007/s00412-002-0212-6.

(43) Calos, M. P.; Lebkowski, J. S.; Botchan, M. R.High Mutation Frequency in DNA Transfected into Mammalian Cells. Proc. Natl. Acad. Sci. U. S. A. 1983. https://doi.org/10.1073/pnas.80.10.3015.

(44) Razzaque, A.; Mizusawa, H.; Seidman, M. M.Rearrangement and Mutagenesis of a Shuttle Vector Plasmid after Passage in Mammalian Cells. Proc. Natl. Acad. Sci. U. S. A. 1983. https://doi.org/10.1073/pnas.80.10.3010.

(45) Stepanenko, A. A.; Heng, H. H.Transient and Stable Vector Transfection: Pitfalls, off-Target Effects, Artifacts. Mutation Research - Reviews in Mutation Research. 2017. https://doi.org/10.1016/j.mrrev.2017.05.002.

(46) Razzaque, A.; Chakrabarti, S.; Joffee, S.; Seidman, M. Mutagenesis of a Shuttle Vector Plasmid in Mammalian Cells. Mol. Cell. Biol. 1984. https://doi.org/10.1128/mcb.4.3.435.

(47) Semenova, N.; Bosnjak, M.; Markelc, B.; Znidar, K.; Cemazar, M.; Heller, L. Multiple Cytosolic DNA Sensors Bind Plasmid DNA after Transfection. Nucleic Acids Res. 2019. https://doi.org/10.1093/nar/gkz768.

(48) Andreeva, L.; Hiller, B.; Kostrewa, D.; Lässig, C.; De Oliveira Mann, C. C.; Jan Drexler, D.; Maiser, A.; Gaidt, M.; Leonhardt, H.; Hornung, V.; Hopfner, K. P.CGAS Senses Long and HMGB/TFAM-Bound U-Turn DNA by Forming Protein-DNA Ladders. Nature 2017. https://doi.org/10.1038/nature23890.

(49) Chen, D.; Murphy, B.; Sung, R.; Bromberg, J. S.Adaptive and Innate Immune Responses to Gene Transfer Vectors: Role of Cytokines and Chemokines in Vector Function. Gene Therapy. 2003. https://doi.org/10.1038/sj.gt.3302031.

(50) Liu, H.; Zhang, H.; Wu, X.; Ma, D.; Wu, J.; Wang, L.; Jiang, Y.; Fei, Y.; Zhu, C.; Tan, R.; Jungblut, P.; Pei, G.; Dorhoi, A.; Yan, Q.; Zhang, F.; Zheng, R.; Liu, S.; Liang, H.; Liu, Z.; Yang, H.; Chen, J.; Wang, P.; Tang, T.; Peng, W.; Hu, Z.; Xu, Z.; Huang, X.; Wang, J.; Li, H.; Zhou, Y.; Liu, F.; Yan, D.; Kaufmann, S. H. E.; Chen, C.; Mao, Z.; Ge, B. Nuclear CGAS Suppresses DNA Repair and Promotes Tumorigenesis. Nature. 2018. https://doi.org/10.1038/s41586-018-0629-6.

(51) Gentili, M.; Lahaye, X.; Nadalin, F.; Nader, G. F. P.; Puig Lombardi, E.; Herve, S.; De Silva, N. S.; Rookhuizen, D. C.; Zueva, E.; Goudot, C.; Maurin, M.; Bochnakian, A.; Amigorena, S.; Piel, M.; Fachinetti, D.; Londoño-Vallejo, A.; Manel, N. The N-Terminal Domain of CGAS Determines Preferential Association with Centromeric DNA and Innate Immune Activation in the Nucleus. Cell Rep. 2019. https://doi.org/10.1016/j.celrep.2019.01.105.

(52) Fenech, M.; Kirsch-Volders, M.; Natarajan, A. T.; Surralles, J.; Crott, J. W.; Parry, J.; Norppa, H.; Eastmond, D. A.; Tucker, J. D.; Thomas, P. Molecular Mechanisms of Micronucleus, Nucleoplasmic Bridge and Nuclear Bud Formation in Mammalian and Human Cells. Mutagenesis. 2011. https://doi.org/10.1093/mutage/geq052.

(53) Umbreit, N. T.; Zhang, C. Z.; Lynch, L. D.; Blaine, L. J.; Cheng, A. M.; Tourdot, R.; Sun, L.; Almubarak, H. F.; Judge, K.; Mitchell, T. J.; Spektor, A.; Pellman, D. Mechanisms Generating Cancer Genome Complexity from a Single Cell Division Error. Science (80-.). 2020. https://doi.org/10.1126/science.aba0712.

(54) Zhang, C. Z.; Spektor, A.; Cornils, H.; Francis, J. M.; Jackson, E. K.; Liu, S.; Meyerson, M.; Pellman, D. Chromothripsis from DNA Damage in Micronuclei. Nature 2015. https://doi.org/10.1038/nature14493.

(55) MacKenzie, K. J.; Carroll, P.; Martin, C. A.; Murina, O.; Fluteau, A.; Simpson, D. J.; Olova, N.; Sutcliffe, H.; Rainger, J. K.; Leitch, A.; Osborn, R. T.; Wheeler, A. P.; Nowotny, M.; Gilbert, N.; Chandra, T.; Reijns, M. A. M.; Jackson, A. P.CGAS Surveillance of Micronuclei Links Genome Instability to Innate Immunity. Nature 2017. https://doi.org/10.1038/nature23449.

(56) Mao, Z.; Bozzella, M.; Seluanov, A.; Gorbunova, V. DNA Repair by Nonhomologous End Joining and Homologous Recombination during Cell Cycle in Human Cells. Cell Cycle 2008. https://doi.org/10.4161/cc.7.18.6679.

(57) Ostrov, N.; Beal, J.; Ellis, T.; Benjamin Gordon, D.; Karas, B. J.; Lee, H. H.; Lenaghan, S. C.; Schloss, J. A.; Stracquadanio, G.; Trefzer, A.; Bader, J. S.; Church, G. M.; Coelho, C. M.; William Efcavitch, J.; Güell, M.; Mitchell, L. A.; Nielsen, A. A. K.; Peck, B.; Smith, A. C.; Neal Stewart, C.; Tekotte, H. Technological Challenges and Milestones for Writing Genomes. Science. 2019. https://doi.org/10.1126/science.aay0339.

(58) Annaluru, N.; Muller, H.; Mitchell, L. A.; Ramalingam, S.; Stracquadanio, G.; Richardson, S. M.; Dymond, J. S.; Kuang, Z.; Scheifele, L. Z.; Cooper, E. M.; Cai, Y.; Zeller, K.; Agmon, N.; Han, J. S.; Hadjithomas, M.; Tullman, J.; Caravelli, K.; Cirelli, K.; Guo, Z.; London, V.; Yeluru, A.; Murugan, S.; Kandavelou, K.; Agier, N.; Fischer, G.; Yang, K.; Martin, J. A.; Bilgel, M.; Bohutski, P.; Boulier, K. M.; Capaldo, B. J.; Chang, J.; Charoen, K.; Choi, W. J.; Deng, P.; DiCarlo, J. E.; Doong, J.; Dunn, J.; Feinberg, J. I.; Fernandez, C.; Floria, C. E.; Gladowski, D.; Hadidi, P.; Ishizuka, I.; Jabbari, J.; Lau, C. Y. L.; Lee, P. A.; Li, S.; Lin, D.; Linder, M. E.; Ling, J.; Liu, J.; Liu, J.; London, M.; Henry, M.; Mao, J.; McDade, J. E.; McMillan, A.; Moore, A. M.; Oh, W. C.; Ouyang, Y.; Patel, R.; Paul, M.; Paulsen, L. C.; Qiu, J.; Rhee, A.; Rubashkin, M. G.; Soh, I. Y.; Sotuyo, N. E.; Srinivas, V.; Suarez, A.; Wong, A.; Wong, R.; Xie, W. R.; Xu, Y.; Yu, A. T.; Koszul, R.; Bader, J. S.; Boeke, J. D.; Chandrasegaran, S. Total Synthesis of a Functional Designer Eukaryotic Chromosome. Science (80-.). 2014. https://doi.org/10.1126/science.1249252.

(59) Richardson, S. M.; Mitchell, L. A.; Stracquadanio, G.; Yang, K.; Dymond, J. S.; DiCarlo, J. E.; Lee, D.; Huang, C. L. V.; Chandrasegaran, S.; Cai, Y.; Boeke, J. D.; Bader, J. S.Design of a Synthetic Yeast Genome. Science (80-.). 2017. https://doi.org/10.1126/science.aaf4557.

(60) Pretorius, I. S.; Boeke, J. D.Yeast 2.0-Connecting the Dots in the Construction of the World’s First Functional Synthetic Eukaryotic Genome. FEMS Yeast Research. 2018. https://doi.org/10.1093/femsyr/foy032.

(61) Pluta, A. F.; Mackay, A. M.; Ainsztein, A. M.; Goldberg, I. G.; Earnshaw, W. C.The Centromere: Hub of Chromosomal Activities. Science (80-.). 1995. https://doi.org/10.1126/science.270.5242.1591.

(62) Earnshaw, W. C.; Migeon, B. R.Three Related Centromere Proteins Are Absent from the Inactive Centromere of a Stable Isodicentric Chromosome. Chromosoma 1985. https://doi.org/10.1007/BF00329812.

(63) Karpen, G. H.; Allshire, R. C.The Case for Epigenetic Effects on Centromere Identity and Function. Trends in Genetics. 1997. https://doi.org/10.1016/S0168-9525(97)01298-5.

(64) Allshire, R. C.; Karpen, G. H.Epigenetic Regulation of Centromeric Chromatin: Old Dogs, New Tricks? Nature Reviews Genetics. 2008. https://doi.org/10.1038/nrg2466.

(65) Ohzeki, J.; Larionov, V.; Earnshaw, W. C.; Masumoto, H. De Novo Formation and Epigenetic Maintenance of Centromere Chromatin. Current Opinion in Cell Biology. 2019. https://doi.org/10.1016/j.ceb.2018.12.004.

(66) Musacchio, A.; Desai, A. A Molecular View of Kinetochore Assembly and Function. Biology. 2017. https://doi.org/10.3390/biology6010005.

(67) Fukagawa, T.; Earnshaw, W. C.The Centromere: Chromatin Foundation for the Kinetochore Machinery. Developmental Cell. 2014. https://doi.org/10.1016/j.devcel.2014.08.016.

(68) Lee, N. C. O.; Larionov, V.; Kouprina, N. Highly Efficient CRISPR/Cas9-Mediated TAR Cloning of Genes and Chromosomal Loci from Complex Genomes in Yeast. Nucleic Acids Res. 2015. https://doi.org/10.1093/nar/gkv112.

(69) Okamoto, Y.; Nakano, M.; Ohzeki, J. I.; Larionov, V.; Masumoto, H. A Minimal CENP-A Core Is Required for Nucleation and Maintenance of a Functional Human Centromere. EMBO J. 2007. https://doi.org/10.1038/sj.emboj.7601584.

(70) Bodor, D. L.; Mata, J. F.; Sergeev, M.; David, A. F.; Salimian, K. J.; Panchenko, T.; Cleveland, D. W.; Black, B. E.; Shah, J. V.; Jansen, L. E. T. The Quantitative Architecture of Centromeric Chromatin. Elife 2014. https://doi.org/10.7554/eLife.02137.

