## Supplementary figures and images for "Analysis of Complex DNA Rearrangements During Early Stages of HAC Formation"

### Supplementary figures

#### Fig. Supplemental 1

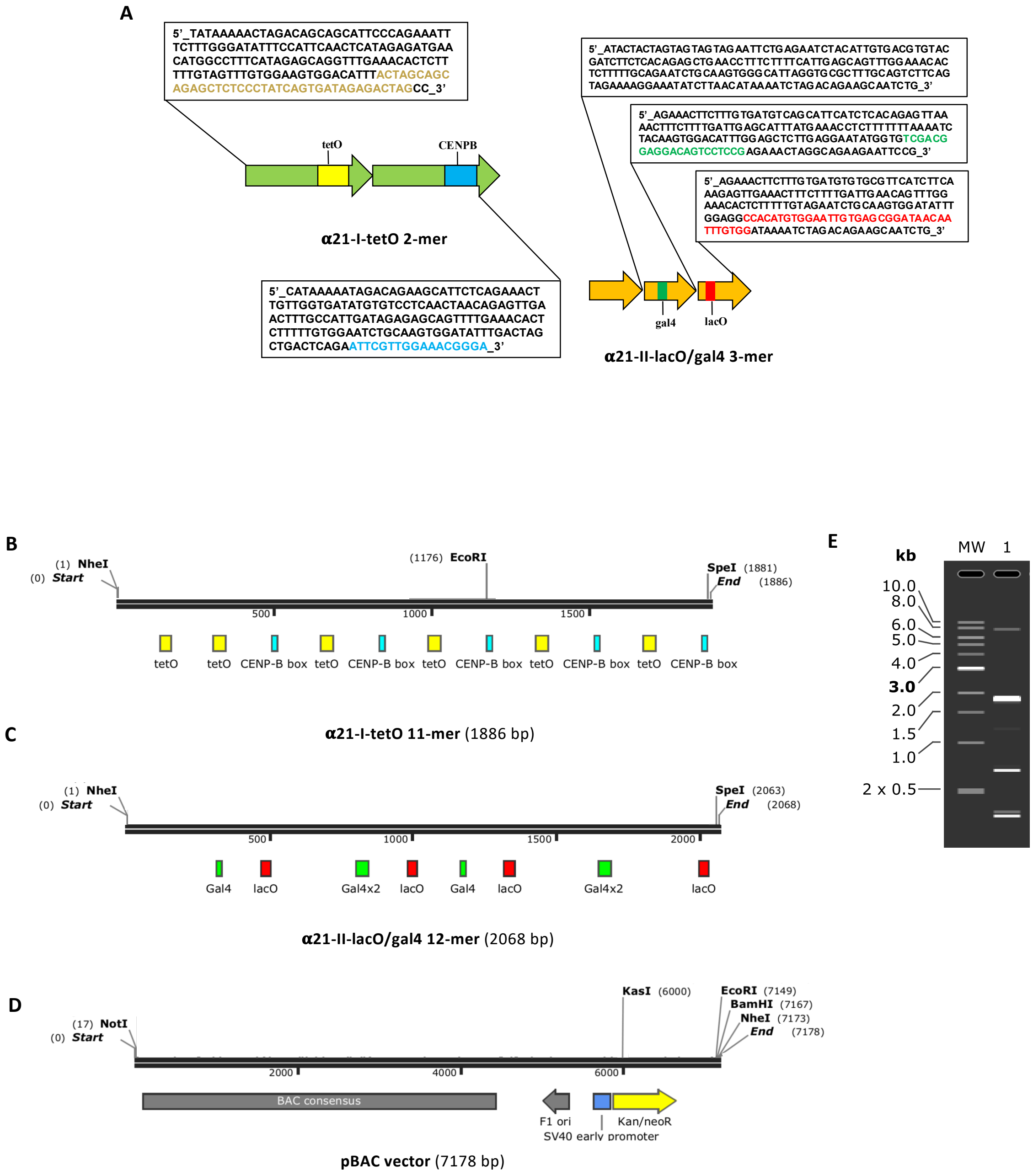

**Fig. Supplemental 2**

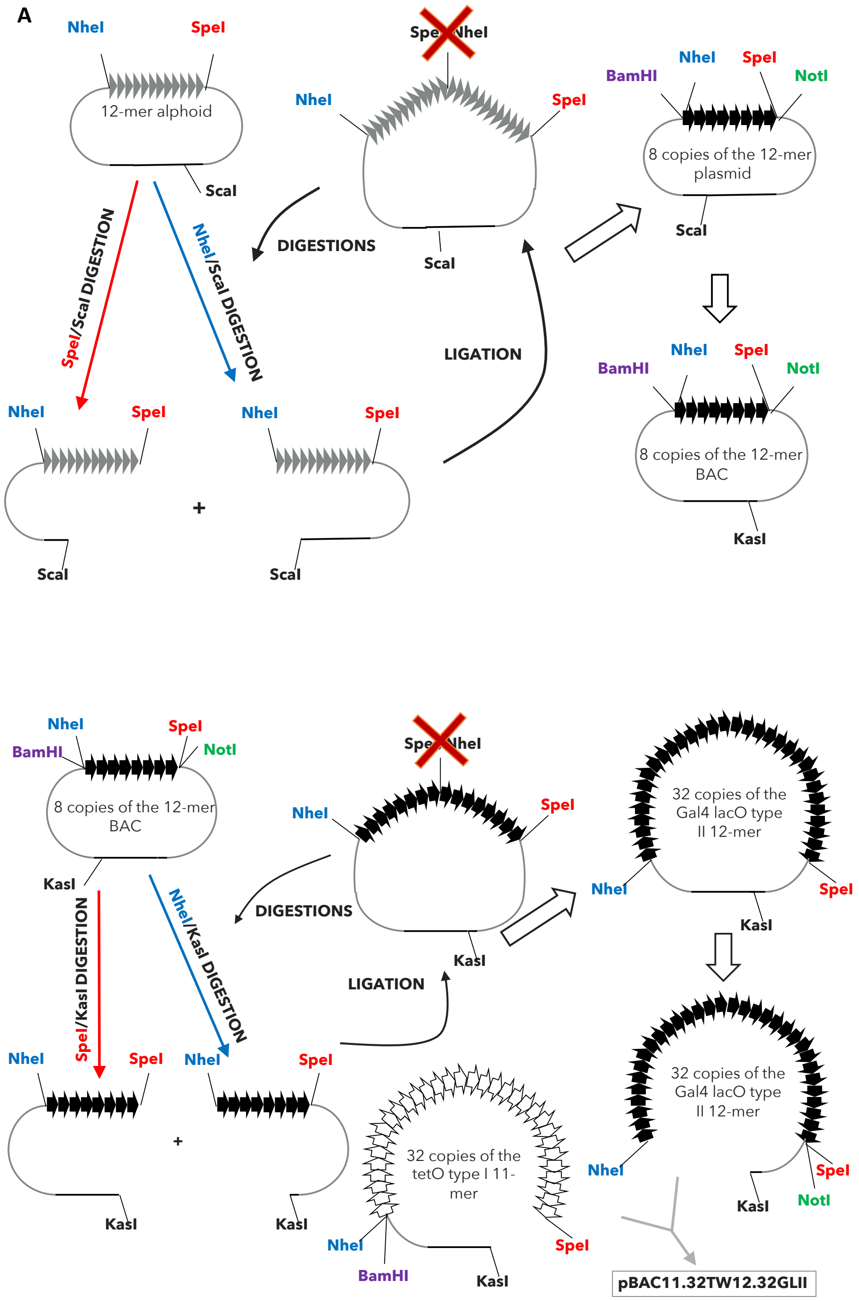

**Fig. Supplemental 3**

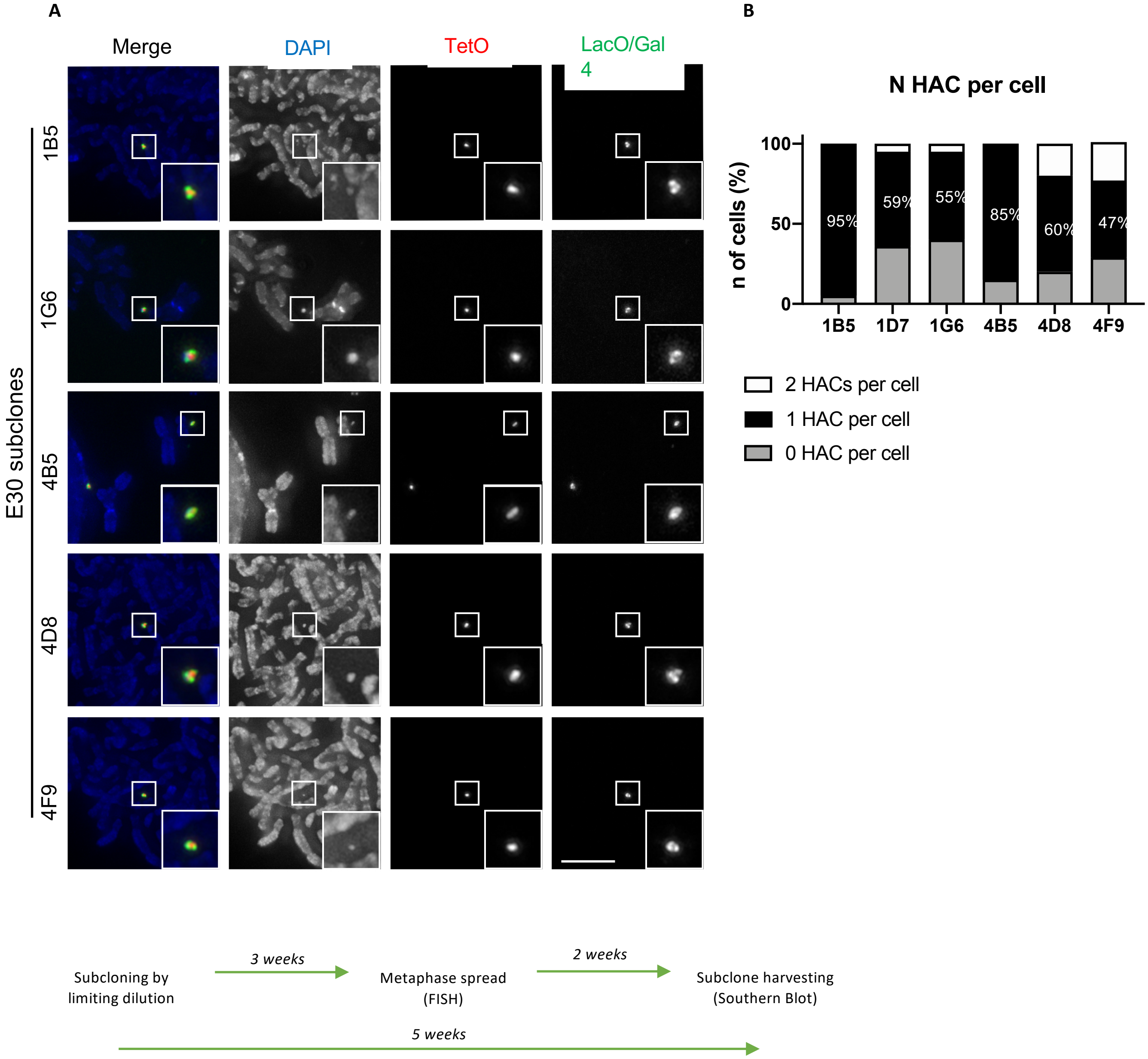

**Fig. Supplemental 4**

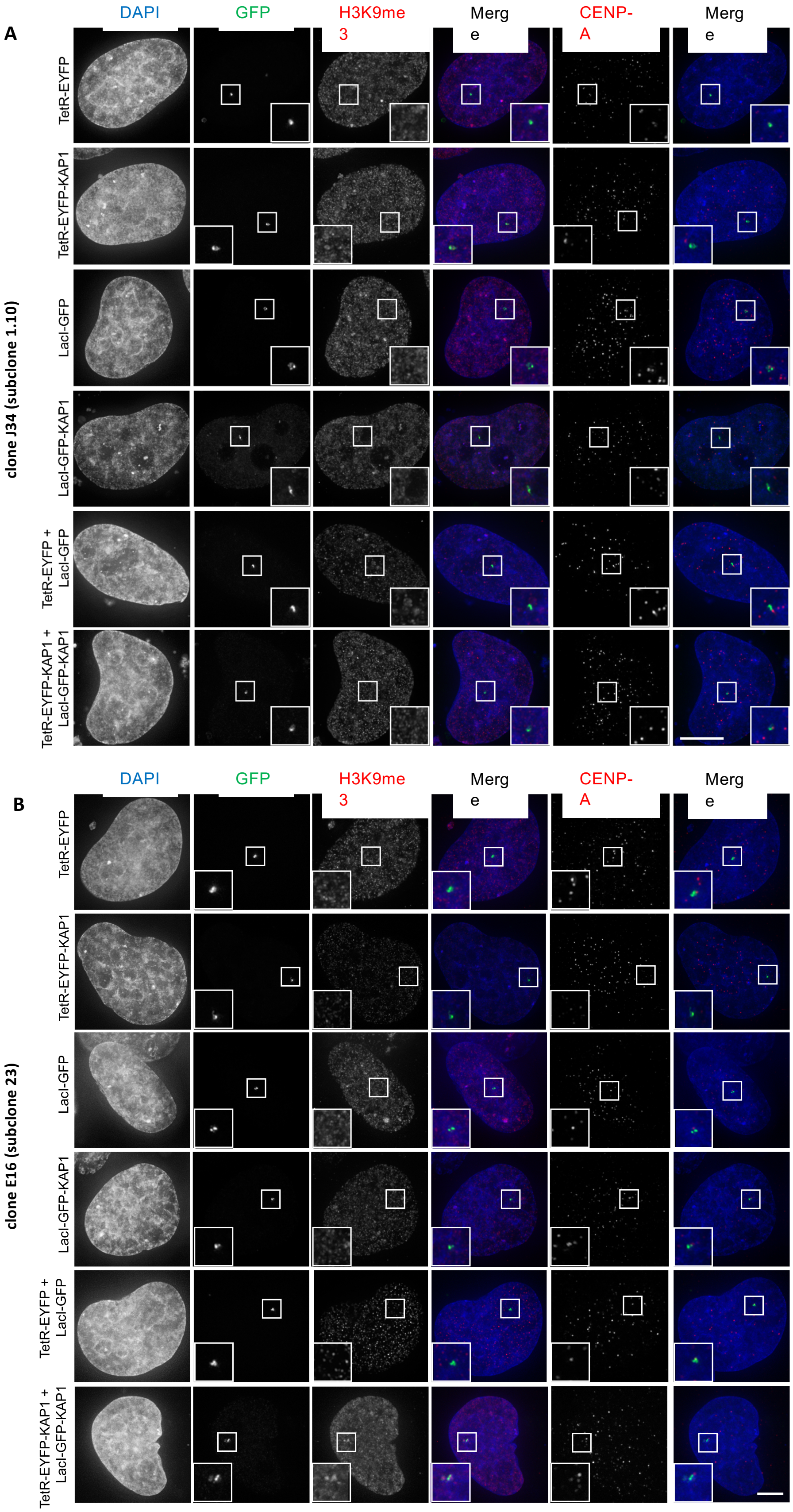

**Fig. Supplemental 5**

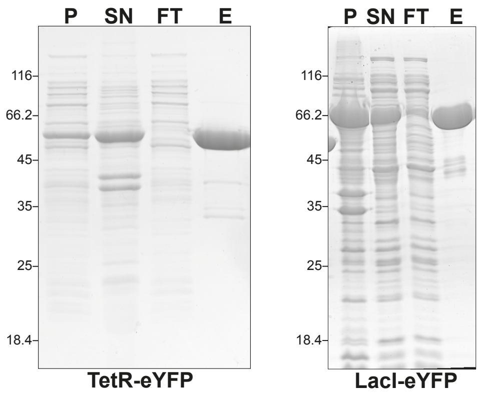
