## Supplementary figure legends for "Analysis of Complex DNA Rearrangements During Early Stages of HAC Formation"

### Supplementary Figures:

Supplementary Figure 1: **Sequences of α21-I$^{TetO}$ array, α21-II$^{LacO/Gal4}$ array and BAC vector.**

(A) Sequences of the α21-I$^{TetO}$ and α21-II$^{LacO/Gal4}$ high ordered repeats (HOR) synthetic DNA, adapted from[4]: both constructs are derived from a chromosome 21 alphoid type I and type II DNA; base pairs in color represent the corresponding feature on the synthetic DNA (colored boxes). (B) Map of α21-I$^{TetO}$ 11-mer (1886 bp) with CENP-B boxes (blue) and TetO (yellow), showing single cut restriction site relevant for the cloning. (C) Map of α21-II$^{LacO/Gal4}$ 12-mer (2068 bp) with LacO (red) and Gal4 (green) sequences, showing single cut restriction site relevant for the cloning. (D) Map of the BAC vector (7178 bp) showing single cut restriction site relevant for the cloning. (E) Pattern predicted in silico using Snapgene of pBAC11.32TW12.32GLII digested with EcoRI (lane 1) run on 1.2% agarose gel. Predicted fragments are 7499, 1880, 677, 370, 342, 340 and 339 bp. (MW 1 kb marker)

Supplementary Figure 2: **Schematic of tandem ligation amplification.**

(A) Multiple cycles of SpeI/ScaI and NheI/ScaI digestions and ligation are performed in plasmid to elongate the synthetic arrays (schematic only showed for α21-II$^{LacO/Gal4}$12-mer, in grey). 8 copies of α21-II$^{LacO/Gal4}$12-mer (in black) are then transferred in BAC. Relevant restriction sites are shown. (B) Multiple cycles of SpeI/KasI and NheI/KasI digestions and ligation are performed to elongate 8 copies of the α21-II$^{LacO/Gal4}$12-mer (in black). 32 copies of the α21-II$^{LacO/Gal4}$12-mer are ligated with 32 copies of the α21-I$^{TetO}$ 11-mer (in white) to form the final pBAC11.32TW12.32GLII. Relevant restriction sites are shown.

Supplementary Figure 3: **Screening of HT1080 subclones by FISH.**

(A) Representative pictures of FISH staining of 6 HT1080 subclones originated from clone E30: slides have been hybridized with DNA probes (TetO-dig/rhodamine α-dig antibody, Gal4-biotin and LacO-biotin/Fitc-streptavidin). DAPI stains DNA. Scalebar = 10µm. Timeline of the experiments performed on HT1080 subclones. (B) Number of metaphases (%) containing 0, 1 or 2 HACs for each subclone; 50 metaphases for each condition were analyzed. Percentages indicate the cells containing 1 HAC.

Supplementary Figure 4: **CENP-A and H3K9me3 have been quantified after transfection with KAP1.** (A, B) Representative images of HT1080 subclones J34 1.10 (A) and E16 23 (B) transfected with the corresponding GFP-targeted protein (green) which localizes the HAC; cells are fixed, permeabilized and stained for α-H3K9me3 rabbit/TRITC α-rabbit and α-CENP-A mouse/CY5 α-mouse antibodies. DAPI stained the nuclei. First merge image from left represent the overlay between DAPI, GFP and H3K9me3; second merge image from left represent the overlay between DAPI, GFP and CENP-A. Scalebar = 10µm.

Supplementary Figure 5: **SDS-PAGE analysis of TetR-eYFP and LacI-eYFP proteins after Ni-NTA affinity purification to assess protein purity.** P= Pellet (insoluble fraction), SN= Supernatant (soluble fraction, input of affinity purification), FT= Flow-Through (unbound fraction), E= Pool of elutions.
